# Identification of Candidate Seed Metabolites and Microbiota Members associated with Germination and Emergence in Common Bean

**DOI:** 10.64898/2026.06.16.732447

**Authors:** Louna Colaert-Sentenac, Elisabeth Planchet, Cyril Abadie, Julie Lalande, Sherif Hamdy, Coralie Marais, Audrey Dupont, Laurence Le Corre, Claude-Emmanuel Koutouan, Marie-Hélène Wagner, Matthieu Barret, Guillaume Tcherkez, Béatrice Teulat, Marie Simonin

## Abstract

**Background:** Seed quality is a complex trait shaped by morphological, biochemical and microbiological properties that are rarely characterised simultaneously, limiting our ability to identify robust predictive indicators of germination speed and seedling emergence across varieties. Here, we performed an exploratory multi-factor characterization of eight common bean (*Phaseolus vulgaris* L.) varieties, combining seed morphometrics, untargeted GC-MS metabolomics on three seed organs, and amplicon sequencing of bacterial and fungal communities, to identify candidate indicators of germination speed and emergence percentage.

**Results:** For all 8 varieties, different seeds from the same lot were used for each analysis. The eight varieties showed substantial variation in both germination speed and seedling emergence traits, used as physiological seed quality proxies. Seed weight and size variation between varieties were correlated with germination speed. The intravariety variance of seed weight was independently correlated with emergence performance. Metabolome composition differed across seed organs, with variety as the dominant driver. Individual-seed metabolomic profiles in the plumule and cotyledon were associated with both germination speed and emergence, yielding each 13 and 14 plumule, and 12 and 2 cotyledon candidate metabolite markers, while hypocotyl-radicle axis yielded only 7 candidate markers associated with germination speed. Fungal community composition was associated with both germination speed and emergence, while bacterial communities were associated with emergence only. Four fungal and seven bacterial taxa were identified as candidate indicators. Inter-kingdom co-occurrence network analysis revealed that fungi with similar germination speed associations tend to cluster in the same modules, suggesting that community-level modules rather than individual taxa may constitute more robust microbial indicators.

**Conclusions:** These results demonstrate that germination speed and emergence capacity are associated with distinct seed properties, and explore morphological, metabolic and microbial candidate indicators and candidate biomarkers to further test for integration into targeted seed quality assessment frameworks for common bean.

## Background

High-quality seeds are essential in agriculture to ensure high germination rate, as well as rapid and uniform emergence and seedling establishment, which affect crop production. Seed quality is a multidimensional property of a seed lot, encompassing genetic identity and purity, physical quality, physiological performance, and seed health status. Seed quality is evaluated through physical criteria (non damaged seeds, uniform size, no insect infestation, moisture content), genetic variety purity, physiological quality (fast and high-rate germination and emergence, protein, oil, fatty acid, and amino-acid content), and health-related criteria (cleanliness, absence of disease-causing microorganisms) [1, 2].

Seed physiological quality is best evaluated by germination and emergence assays, conducted with precise parameter settings for official evaluation [3]. Germination covers the resumption of metabolic activity and embryo growth after imbibition up to the radicle protrusion, while seedling emergence covers the following elongation of the seedling and penetration through the substrate up to the surface. However, germination and emergence assays are time-, resource-, and material-consuming. To evaluate seed physiological quality, morphological proxies are historically the most commonly used indicators, with many studies linking seed weight and seed energy stock quantity, seed color and imbibition time before germination [4]. During the transition from quiescent dry state to active growth, seeds rely on tightly regulated metabolic networks involving carbohydrates, lipids, amino acids, antioxidants, and signaling molecules that support energy supply, stress tolerance, and developmental progression [5, 6]. Consistent with this view, seed metabolic signature could provide valuable insight into seed germination ability, as previously evidenced for maize, tomato, rice, and wheat [7–9]. Indeed, as evidenced for dry seed mRNAs and proteins [10–12], metabolome could constitute a reservoir of ready-to-use molecules when the imbibition starts, thus impacting seed germination. Metabolic pathways involved in seed quality and seed germination are traditionally explored through genome-wide association study (GWAS), transcriptomic, or proteomic studies, but the dry seed metabolite pool has yet received little attention. The difference between the composition of seed organs could be especially relevant, given their specialized roles in germination [13]. In parallel, the seed microbiota is increasingly viewed as a functional component of seed biology rather than a passive contaminant. Seeds carry microbial communities that can contribute to plant establishment, health, and resilience [14–16], and are now considered a primary inoculum for the next plant generation. Yet routine seed quality evaluation usually focuses on harmful microorganisms and does not account for potentially beneficial seed-associated taxa. As a result, the relationships between seed microbiota, seed metabolome, and germination and emergence performance remain poorly resolved, as well as their contribution to the differences in seed quality between varieties, with only few studies exploring their interactions [17].

Here, we addressed this issue with a comprehensive multi-factor study to explore traits associated with common bean (*Phaseolus vulgaris* L.) seed germination and seedling emergence, taking advantage of a collection of French varieties of commercial significance. Common bean was selected as model legume species due to its importance as staple food, displaying high genetic and phenotypic diversity [18, 19]. As one of the most widely cultivated grain legumes, common bean provides essential protein amino acids, fibers, and micronutrients, making it a critical component of human diets, especially in developing countries [19–21]. In addition, bean varieties exhibit substantial diversity in seed morphology, composition, and germination behaviour, offering an excellent opportunity to explore correlations between these traits and overall seed quality [22], that to the best of our knowledge has not yet been studied. Seeds used for each analysis were subsampled from batches of seeds, obtained from plants cultivated at the same time in the same location so as to limit environmental effect on seed development and microbiota assembly.

Germination and emergence assays, GC-MS metabolomic analyses of various seed tissues, and analyses of the seed microbiome across varieties were used to identify candidate indicators of germination and emergence. We examined their correlation with seed properties using multivariate statistical methods. Our working questions were: (i) how do mature seed morphological, metabolic, and microbiota-related parameters relate to seed germination speed and seedling emergence percentage, and can we therefore identify candidate quality indicators and biomarkers? (ii) are such candidate indicators and biomarkers valid across different bean varieties?

## Methods

All statistical analyses were conducted on R Statistical Software (v.4.5.3; R Core Team 2025). All figures were generated using *ggplot2* v4.0.2 package. A list of package versions used is provided in the code.

### Plant material

Eight *Phaseolus vulgaris* L. (common bean) varieties were selected by the seed company Vilmorin-Mikado SAS (Limagrain group) to represent a large panel of cultivated bean varieties in France (Caprice, Contender, Deezer, Facila, Flavert, Linex, Vanilla, and Vezer, **Figure S1A**), including dwarf French string bean, and shelling bean. The seeds for all varieties were produced in 2020 as part of a single field trial from plants grown from commercial seeds (Vilmorin-Mikado SAS), at the National Federation of Seed Multipliers experimental station (FNAMS, 43°57’35.654”N 0°23’45.428”E, Condom, Gers, France, **Figure S1B**). Culture was done under the following farming conditions: The herbicide Mercantor Gold (1.4 L/ha) was applied once 2 days before sowing. The grass herbicide Stratos® Ultra was applied once one month after sowing at 4 L/ha, and the field was irrigated regularly. The culture was conducted in a single plot separated into 5 x 2 bands of soil (five bands of 2.8 m wide spaced by a 0.55 m space, on two rows of 25 m long separated by a 4 m space). Each variety was sown in a separate band to ensure the purity of the harvested seeds at a density of 30 plants per m^2^, with the variety band randomized across the field. Harvested seeds were dried at room temperature, cleaned by filtering soil and non-seed structures on sieves at the National Seed Testing Station (GEVES, France), and stored in paper bags at 4°C in the dark until further use.

### Seed germination and seedling emergence indicators

#### Seed germination assays

Seed germination assays were performed at the National Seed Testing Station (GEVES, France) on wet pleated paper in a Jacobsen germinator set at 20°C, 95% humidity and equipped with four RGB cameras [23]. The assay was performed on 4 biological replicates of 24 non-damaged seeds for each variety, sown in different boxes. A seed was considered germinated as soon as the radicle emerged from seed testa. Germination imaging was recorded every 2 h over 6 days (144 h). Germination time-course curves for each variety were used to determine two germination indicators: the final germination percentage (FGP) and its variety average (mean FGP), and the germination speed per replicate (time to 50% of germinated seeds - t50germ) and its variety average - mean t50germ).

#### Seedling emergence assays

Seedling emergence assays were performed on the Phenotic platform [24] on six biological replicates of 20 seeds without defaults for each variety, sown in the same tray. Seeds were grown in potting mix [K® TraySUBSTRAT Klasmann-Deilmann France] in seedling trays, with tray position randomised across varieties. Plants were grown in a climatic chamber set at 23°C day and 20°C night, for 16 h day and 8 h night, under 70% humidity and 150 µmol·m^-²^·s^-1^ LED light (**Figure S1C**). The number of non-germinated seeds, normal seedlings that have emerged, or abnormal seedlings was recorded on the seventh day according to International Seed Testing Association (ISTA) guidelines for common bean [3]. Final emergence percentage (FEP) and its variety average (mean FEP) were used as the indicator of emergence of varieties.

#### Statistical analyses - Germination and emergence indicators

Germination kinetic curves were fitted per variety using a four-parameter Hill function (FourPHFfit.bulk, germinationmetrics [25]) to derive the time to 50% of germinated seeds for each replicate (t50germ) [26]. FGP, t50germ and FEP were compared between varieties by one-way ANOVA (linear model fitted with the lm function, Type II ANOVA tested with car::Anova) after verification of residual normality (Q-Q plot and Shapiro-Wilk test) and homoscedasticity assessment (diagnostic plots). Post-hoc pairwise comparison between variety means were performed using estimated marginal means with Tukey correction (emmans), and letter display groups were assigned at α = 0.05 (multcomp package).

### Seed weight and morphological characteristics

#### Seed measurements

Seed morphological characteristics were measured on 100 individual seeds, considered as biological replicates, for each variety at GEVES (France). Their morphological characteristics, including seed area (**Table S1**), were determined from images acquired by 2D X-rays radiography with digital X-ray equipment Faxitron MX-20 (Faxitron X-ray Corp., Wheeling, IL, U.S.A.) [23]. These images were analysed using Python Programming Language, OpenCV, and AVIZO (Thermo Fisher Scientific). This approach also enabled the identification of seeds presenting mechanical defects (visual inspection for tegument fissures, protruding teguments, insect damage and spread cotyledons). The proportion of seeds presenting defects was quite low (0-16.7% across varieties, 3.8% overall), and those seeds were excluded from further processing. All seeds were individually weighed (mg) on a precision balance.

#### Statistical analyses of seed measurements

To assess variations in seed weight and size diversity between varieties and their associations with t50germ and FEP, univariate statistics were used.

Variety-level means and standard deviations were computed for seed weight and all morphological variables across the 100 seeds per variety. Thanks to individual measures, the coefficient of variation (CV = SD/mean) was also computed for seed weight and area as an index of intravariety size heterogeneity. Between-variety Spearman rank correlations were computed between variety-level means of all morphological variables and mean t50germ and mean FEP, using the eight variety means as the unit of analysis (n = 8). Significance was assessed at α = 0.05 without and with BH correction for multiple testing given the exploratory nature of the analysis.

The relationship between individual seed weight and projected area was visualised using scatter plots with variety-level 95% confidence ellipses. Logarithmic regression (y ∼ log(x)) was used to model the relationship between mean seed weight and mean t50germ across varieties, as visual inspection suggested a non-linear relationship. Linear regression was used for all other trait pairs. *R*² values from ordinary least squares regression are reported alongside Spearman rank correlation coefficients to characterise both the strength and the functional form of the relationships.

To test whether the between-variety relationship between seed size and germination speed was recapitulated at the individual seed level, Spearman rank correlations between individual seed weight or area, and individual germination time from the corresponding flat paper germination assay were computed separately within each variety, restricted to germinated seeds (n = 43-90 per variety depending on germination rate).

### Seed metabolome analysis

#### Sample preparation, extraction, and GC-MS analysis

The metabolomes of different seed organs (plumule, cotyledons, and hypocotyl-radicle - HR-axis) were studied by GC-MS across all varieties. For each variety, four mature dry seeds of small, medium, and big seed size relative to the variety mean (12 seeds per variety) were dissected to isolate the organs. Each organ was weighted and ground into a fine powder with metallic beads. Extraction of metabolites were carried out on 4 mg dry weight with 200 µL 90%/10%(v/v) methanol - MilliQ water with adonitol (55 µmol L^-1^, Sigma-Aldrich®) as an internal standard. The extraction volume was adjusted proportionally for samples < 4 mg (plumule, HR-axis). Extracts were centrifuged for 10 min at 1,500 *g*. For each extract, 10 µL of supernatant was transferred into a vial with insert, and 10 µL was used to prepare a quality control (QC), i.e. a mix of all seed samples, which was also prepared in 10 µL per vial. All vials were spin-dried overnight at room temperature, sealed and kept in dark dry conditions until GC-MS analysis.

GC-MS analyses were conducted with a GC-MS-Orbitrap Q Exactive (Thermo Scientific). Samples were automatically derivatized with 20 µL of 20 mg·mL⁻¹ methoxyamine-pyridine solution for 90 min at 37°C, then with 30 µL of N-methyl-N-(trimethylsilyl)-trifluoroacetamide (MSTFA) for 30 min at 37°C. Five µL of alkane mix (C_9_ to C_36_ alkanes at 3 µg·µL⁻¹, Connecticut n-Hydrocarbon Mix, Supelco) was added before injection to compute the retention index. One µL sample was injected in splitless mode in a TG-5 SILMS column (30 m x 0.25 mm x 0.25 µm, Thermo Scientific) at 230°C, using a Trace 1300 Series GC (Thermo Scientific). Samples were carried with a constant helium flow (1 mL·min⁻¹). Temperature in the GC oven was set at 70°C for 1 min, then raised at 15°C·min⁻¹ to 325°C and kept such for 4 min.

MS analyses were done in positive polarity in full MS scan mode with source set at weight scan range: 50-750 m/z, resolution: 60,000, AGC: target 1E6, MS transfer line: 300°C, and filament delay: 4.12 min. Metabolites were ionized by electron impact at 70 eV, 250°C. Blanks were injected at the beginning and at the end of each batch to assess column cleanliness.

QC samples were added to each batch to assess system stability throughout the batches and to condition the analytical platform. Analytes were identified automatically with TraceFinder® (Thermo Scientific) using retention time and m/z ion of major characteristic fragments and confirmation fragment(s) by comparison to a local compound database [27]. Peak integration by TraceFinder® was checked manually for all metabolites. To avoid too much bias in metabolome composition due to seed size, only seeds with a weight within ± 50% standard deviation around the mean seed weight for each variety were kept [28]. Integral values were normalized using the internal standard.

#### Statistical analysis of metabolomic data

Relative analyte abundances were imported in R. A total of 252 annotated analytes were identified across organs. Metabolites detected in less than 20% of samples within an organ were excluded, leading to 221 retained metabolites per organ. Data were Z-scaled. Outlier samples were identified by principal component analysis (PCA) and removed when outside the Hotelling’s T² ellipse at the 95% confidence interval. As a result, 38 cotyledon, 53 HR-axis, and 46 plumule samples across 8 varieties were kept, and used as biological replicates. Due to the filtering steps, the per-seed association was kept for 31 seeds with all organ samples remaining, 20 seeds with 2 organ samples, and 4 seeds had only one organ sample remaining.

Global metabolome variation across organs was explored by PCA on data Z-scaled across all samples jointly (**Table S5**). A heatmap was generated from these filtered, Z-scaled data using ComplexHeatmap v.2.28.0 [29]. Metabolic differences between organs were assessed by PERMANOVA (adonis2, vegan package v2.6, Euclidean distance, 999 permutations). Homogeneity of multivariate dispersion across organs was tested with betadisper followed by permutest (999 permutations, pairwise comparisons). Metabolites with significantly higher median abundance in a given organ relative to at least another organ were identified using linear mixed models with organ as a fixed effect and individual seed identity as a random effect to account for the matched dissection design (lmer function). For metabolites where the random effect variance was estimated at zero (singular fit), a standard linear model was used as a conservative fallback (lm function). P-values for the organ fixed effect were extracted using Type III ANOVA with Satterthwaite’s method (lmerTest package) and adjusted for multiple testing with Benjamini-Hochberg (BH) correction (adjusted p < 0.05), followed by Dunn post-hoc pairwise comparisons. Per-sample normalized abundances of the metabolites enriched in at least one organ were reported in **Table S6**.

For each organ, PCA was performed on each per-organ Z-scaled metabolome (**Table S7**). Individual seed weight, organ-specific seed weight, mean t50germ and mean FEP were projected on the per-organ PCA space using their Pearson correlation coefficients with PC1 and PC2 scores as vector coordinates, scaled by a factor 10 for visualization. Arrow length and direction therefore reflect the strength and direction of the linear association between each metadata variable and the metabolomic principal component axes. PERMANOVA statistics (R² and p-value) testing the association of variety, mean t50germ and mean FEP with metabolome composition were computed using the vegan::adonis2 function on Euclidean distances of per-organ Z-scaled profiles.

The association between metabolome and t50germ or FEP was assessed by orthogonal partial least squares regression (OPLS, ropls v.1.44.0) [30], with metabolites as X and per-replicate t50germ or FEP as continuous Y. Models used one predictive component; orthogonal components were determined automatically (median 2, range 1–6). Between 214 and 220 metabolites entered the models, for 38 (cotyledon), 46 (plumule) and 53 (hypocotyl-radicle axis) seeds from eight varieties. Because metabolome analysis is destructive, the same seeds could not be used for metabolome analysis and for germination and emergence assays. To propagate the uncertainty of the t50germ- or FEP-metabolome pairing, OPLS was run on 100 independent random within-variety permutations, each pairing metabolome samples of one organ with per-replicate t50germ or FEP values measured on seeds of the same lot and variety; this number was set by a preliminary stability analysis confirming convergence of significance frequencies and of loading sign. Model quality across permutations is reported in **Table S9**. Three complementary tests were used to characterise model performance. First, the ropls response permutation test (200 permutations) permutes Y across all samples irrespective of variety and tests whether any association exists between X and Y. All models fell below the resolution limit of this test. Second, because a variance-components model (t50germ ∼ 1 + (1|Variety)) attributed 94.1–96.2% of t50germ variance to variety, Y is close to constant within a variety and models fitted under cross-validation folds drawn at random over individual seeds may recover it by recognising the variety rather than by relating the metabolome to the phenotype. To test this, variety mean phenotypes were permuted among the eight varieties and reassigned to their seeds, and Q² was recomputed under the same 7-fold scheme (200 permutations; 8! = 40,320 distinct assignments exist). This null preserves the variety structure of the data and differs from the observed data only in which phenotype is attached to which variety. Q² was compared with this null distribution, and against the similarity between permuted and observed phenotypes. Third, leave-one-variety-out (LOVO) cross-validation held each variety in turn, with centring, unit-variance scaling and the zero-variance filter refitted on the training varieties only and complexity fixed a priori at one predictive and one orthogonal component. Significance was assessed against shuffling phenotypes across the eight varieties (199 permutations). The OPLS was therefore presented as a candidate-discovery procedure rather than a predictive model.

A metabolite was retained as a candidate if it met simultaneously: (i) large loading value in the OPLS, i.e. |pq1| > 0.05, and its variable projection VIP > 1 in ≥ 70% of permutations; (ii) univariate significance i.e. Spearman correlation with t50germ or FEP with Benjamini-Hochberg-adjusted p < 0.05 in ≥ 70% of permutations; and (iii) the sign of the OPLS loading pq1 was consistent across ≥ 80% of permutations. Results were visualized as volcano plots (mean pq1 vs. −log₁₀ mean BH-adjusted p-value). Stability of the loading and of the significance across permutations is reported for each candidate (CV of pq1 and of −log₁₀ BH-adjusted p; **Table S10**). This analysis identified an initial set of candidate metabolites. Candidates were then examined by two complementary approaches, both independent of the within-variety permutation : (i) Pearson or Spearman correlations (n = 8 varieties, test selected based on Shapiro-Wilk normality assessment) between variety-median scaled metabolite abundances and variety mean t50germ or FEP. Because the number of varieties (n = 8) limits the power of individual tests and no correlation survived Benjamini–Hochberg correction across the 214–220 metabolites tested, a family-level permutation test was performed: variety mean phenotypes were shuffled across the eight varieties (9,999 permutations, Spearman) and the number of metabolites exceeding |ρ| thresholds was recounted. In no organ × variable combination did the observed count exceed the permuted distribution (p = 0.19-0.94, **Table S10**). (ii) Elastic net regression with cross-validation folds defined by variety (glmnet v.5.0, [31], α optimised by cross-validation), performing independent sparse variable selection from the full metabolome, thus testing whether OPLS-identified candidates are also among the most parsimonious predictors of t50germ or FEP when all metabolites compete simultaneously for inclusion in a penalised regression model (applied on individual samples). The normalized abundance of retained candidate biomarkers is reported in **Table S10**.

### Seed microbiota analysis

#### Seed preparation, DNA extraction, library construction and sequencing

Due to the very low biomass of the microbial communities per individual bean seed [32], seed microbiota analyses were conducted by metabarcoding on three biological replicates of macerate from batches of 30 seeds each, for each variety.

Seed batches were rinsed three times with sterile water to remove dust or debris on the surface of the seeds. Seeds were then soaked in 2 mL per gram of seeds of Phosphate Buffer Saline sterile solution (Sigma-Aldrich, Saint-Louis, MO, USA) supplemented with 0.05% [v/v] of Tween 20 (Sigma-Aldrich), under agitation (220 rpm) at 6°C for 16 h, following the ISTA guidelines to collect both epiphytic and endophytic microbiota member. The liquid obtained after seed soaking was centrifugated in 15 mL tubes (4,000 *g*, 15 min, 20°C), pellet was resuspended in 1 mL surnatant and transferred in a 1.5 mL tube, and stored at −20°C until extraction. DNA was extracted from the resuspended pellet using NucleoSpin ® 96 Food kit (Macherey-Nagel, Düren, Germany), following the manufacturer’s instructions. Bacterial and fungal DNA, along with extraction blank and controls, were amplified through PCRs with a high-fidelity Taq DNA polymerase (AccuPrime Taq DNA Polymerase System, Invitrogen, Carlsbad, CA, USA), with primers for a portion of the bacterial *gyrB* gene (gyrB_aF64/gyrB_aR353) [33] and primers for region 1 of the fungal internal transcribed spacer of nuclear DNA (ITS1F/ITS2) [34], respectively. A blank extraction kit control, a PCR-negative control, and a PCR-positive control (*Lactococcus piscium* DSM6634, a fish pathogen that is not plant-associated) were included in each PCR plate. The first PCR was performed with 5 µL of 10X buffer, 1 µL of forward and reverse primers (100 µM for *gyrB* and 10 µM for ITS), 0.2 µL of Taq and 5 µL of DNA for a final volume of 50 µL per sample. Cycling conditions were such : initial denaturation step at 94°C (3 min), then 35 cycles of amplification at 94°C (30 s), 55°C for *gyrB* and 50°C for ITS (45 s), and 68°C (90 s), then a final elongation step at 68°C (10 min). Amplicons were purified with magnetic beads (Sera-MagTM, Merck, Kenilworth, New Jersey, USA). A second PCR was performed to incorporate Illumina adapters and barcodes, with 5 µL of 10X buffer, 0.2 µL of Taq, and 10 µL of PCR1 amplicons and 2 µL of indexes for a final volume of 50 µL per sample. Cycling conditions were such: initial denaturation step at 94°C (1 min), then 12 cycles of amplification at 94°C (1 min), 55°C (1 min), and 68°C (1 min), then a final elongation step at 68°C (10 min). Amplicons were again purified with magnetic beads, quantified with QuantIT PicoGreen dsDNA Assay Kit (Invitrogen) and pooled in equimolar concentration. Concentration of the pool was measured via quantitative PCR (KAPA Library Quantification Kit, Roche, Basel, Switzerland). Amplicon libraries were mixed with 5% PhiX and sequenced with MiSeq reagent kits v3 600 cycles (Illumina, San Diego, California, USA) for 2 x 250 bp on the Illumina MiSeq platform (ANAN platform, SFR QuaSav, Angers, France). The raw amplicon sequencing data are available on the European Nucleotide Archive (ENA) with the accession number PRJEB59579 (files named V[variety number]S1, with variety number from 1 to 8: Deezer, Caprice, Linex, Vanilla, Facila, Vezer, Contender, Flavert).

#### Sequencing data processing

Sequenced read integrity was verified for file integrity with gzip, then processed using QIIME2 v2019.7 [35]. Primer trimming and read demultiplexing were performed with *cutadapt 2.7* plugin. Denoising (including merging for *gyrB* marker), chimera filtering and Amplicon Sequence Variants (ASVs) inference were conducted using the DADA2 (v1.10.0) [36] plugin within QIIME2 with max-ee = 2. For the *gyrB* gene, truncation parameters were set to 180 bp (forward) and 160 bp (reverse). For the ITS marker, the single-end mode using the forward reads only was used without truncation nor trimming as recommended by [34]. ASV taxonomy was assigned using a feature-classifier based on Naïve Bayes algorithm [37, 38] based on the UNITE database for ITS1 (UNITEv10 (tgz), version 19.02.2025) [39]; and on our local *gyrB* database (available in ref [40] under train_set_gyrB_v5[40], with default bootstrap confidence threshold, minBoot = 50, tryRC = FALSE. For marker *gyrB*, ASVs assigned to “parE” (a *gyrB* paralog) or unclassified at the phylum level were excluded from further analysis. For marker ITS, ASVs not classified as Fungi (e.g., Viridiplantae, Protista) or unassigned were excluded. The resulting feature tables were filtered to retain ASVs present in at least two samples with a minimum number of 20 reads. Samples with fewer than 1000 total reads were excluded to mitigate bias due to low sequencing depth. ASV tables, representative sequences, and associated taxonomic classifications were exported to R for downstream ecological analysis. ASVs were manually decontaminated by filtering ASVs present in the two PCR-positive controls, ASVs present in less than three seed batch samples, and ASVs present in at least three PCR-negative controls. A total of 20 ASVs were removed from the gyrB gene marker amplification and sequencing, and 11 ASVs from the ITS marker amplification sequencing.

#### Statistical analyses of microbial community composition

Microbiota analyses were conducted with the Phyloseq package v1.54.2 [41]. Alpha rarefaction curves were generated to assess sequencing depth sufficiency. For alpha diversity metric calculations, the gyrB dataset was rarefied using coverage-based rarefaction following Hsieh et al recommendations [40] (target sample coverage of 99.98%, corresponding to a mean depth of 8,975 reads per sample), while the ITS dataset was rarefied to an even depth of 7,625 reads per sample, corresponding to the minimum observed sequencing depth across samples, as coverage-based rarefaction was not applicable due to the absence of singletons. α-diversity (richness, Chao1, Shannon) was estimated on rarefied data and differences between varieties tested with Kruskal-Wallis tests followed by Dunn’s post-hoc comparisons with Benjamini-Hochberg correction. β-diversity was assessed on unrarefied data filtered at 5% prevalence and transformed with robust centered-log ratio, using PCoA on Euclidean distances for visualisation and PERMANOVA (999 permutations, vegan::adonis2) for statistical testing, with homogeneity of dispersions verified beforehand using vegan::betadisper. Taxonomic composition was visualised at genus level from non-rarefied counts filtered at 5% prevalence, with genera below 1% maximum relative abundance grouped as ‘Other’. Those classical microbial analyses, along with their results, are detailed in **Note S1**.

Associations between variety-mean germination and emergence (mean t50germ, meanFEP) and seed microbiota composition were assessed for bacterial (*gyrB*) and fungal (ITS) communities separately, on the non-rarefied read counts filtered by prevalence ≥ 15% to avoid taxa present only in replicates of one variety. The relation between community composition and t50germ, or FEP, was tested with PERMANOVA on the rCLR-transformed Euclidean distance matrix with 999 permutations. As mean t50germ and mean FEP were nearly uncorrelated (Spearman r = −0.05) and showed no collinearity (VIF < 2 for both), the relation between microbiota and each variable was tested in separate models to maximize statistical power given the limited sample size (n = 21). Because mean t50germ and mean FEP were each defined as a single value per variety, these traits are structurally confounded with variety identity; their effects on community composition could therefore not be statistically disentangled from a variety effect. The reported associations should accordingly be interpreted as reflecting covariation between variety identity, germination phenotype, and microbiota composition. Associations were visualized using distance-based redundancy analysis (db-RDA, vegan::dbrda), with species scores assigned from the rCLR matrix (vegan::sppscores). The ten ASVs contributing most to the constrained axes were identified by their Euclidean distance from the origin in ordination space and labeled using the best available taxonomic assignment.

As for metabolomic data, seeds used for microbiota analyses were distinct (albeit from the same lot) as those used for germination and emergence assays. To identify individual ASVs whose relative abundance was associated with t50germ and FEP, five complementary statistical approaches were applied to non-rarefied read counts filtered by prevalence (≥ 15% of samples):

i. Variety-level Spearman correlation (SpearmanVar) with BH correction, on per-variety means of microbial relative TSS-normalized abundances (n = 8 varieties), and mean t50germ or mean FEP values, tested if the correlation held at the variety level,
ii. Within-variety permutation Spearman correlation (SpearmanPerm) with BH correction, on 100 random within-variety permutations of microbiota data and t50germ or FEP values (n = 24 samples across 8 varieties, same permutation design as for metabolomic data: random association between per-replicate t50germ or FEP values with microbiota samples of the same variety), tested if samples with higher relative abundance of this taxon tend to have lower t50germ and higher FEP. A taxon was retained as a candidate if it met simultaneously: BH-adjusted q < 0.25 in ≥ 70% of permutations, and a consistent sign of association in ≥ 80% of permutations. This approach mirrors the permutation strategy applied in the metabolomics analysis.
iii. Linear model-based association analysis (MaAsLin2, Maaslin2 R package, TSS normalization, log transformation, Benjamini-Hochberg correction, using the same within-variety permutation design), tested if there was a linear association between the log-transformed relative abundance of a taxon and t50germ or FEP (n = 24 samples, same permutation design), after accounting for compositionality through normalisation.
iv. Compositional correlation analysis (ALDEx2 v1.44.0, Monte Carlo Dirichlet sampling, 128 instances, Spearman correlation test), applied to individual sample-level raw counts (n = 24 samples) tested if the association between a taxon and the variety-mean t50germ or FEP value was robust to the stochastic uncertainty inherent in compositional count data, when abundance is expressed relative to the geometric mean of all taxa.
v. Finally, bias-corrected compositional differential abundance analysis (ANCOMBC2, ANCOMBC R package) applied to the same individual sample-level counts (n = 24), tested if the association between a taxon and variety-mean t50germ or FEP was robust to differences in total microbial load between samples.

A relaxed significance threshold of q ≤ 0.25 was applied across all methods given the limited effective sample size (n = 8 varieties), and results are interpreted as exploratory. The multiplicity of methods helped strengthen confidence in candidate ASVs that were flagged across statistical frameworks. However, because all five approaches were applied to the same dataset, they should not be regarded as independent biological validations. Candidate ASVs identified here should therefore be interpreted as exploratory hypotheses requiring confirmation in an independent, adequately replicated dataset. Cross-method consistency was assessed by building a comparison table recording, for each taxon-variable combination, which methods identified a significant association and the corresponding effect sizes (Spearman rho from SpearmanVar and SpearmanPerm, MaAsLin2 log fold-change, ALDEx2 Spearman rho, ANCOM-BC2 log fold-change). Taxa confirmed by at least two independent methods were considered robust candidates and retained for further visualization.

To better understand the microbiota structure, a co-occurrence network approach was used. Analyses were performed on non-rarefied read counts filtered by prevalence (≥ 15% of samples) for both bacterial (*gyrB*, n = 96 ASVs) and fungal (ITS, n = 93 ASVs) communities. The co-occurrence of bacterial and fungal taxa was inferred with a sparse co-abundance network, using the SPIEC-EASI algorithm - SpiecEasi R package, v1.99.0 [42] - that estimates sparse inverse covariance matrices from compositional microbiome data using Meinshausen-Bühlmann neighbourhood selection method fitting sparse regression models for each node against all others. For multi-kingdom analysis, we used the multi.spiec.easi function, which handles compositionality within each domain separately while estimating cross-domain associations jointly. Network sparsity was optimized using the StARS (Stability Approach to Regularization Selection) criterion on 20 values of λ (minimum ration: 10^-2^), with 50 subsampling repetitions, a stability threshold of 0.05, and a subsampling ratio of 0.8. The optimal regularization parameter was selected at the lambda value first exceeding the StARS stability threshold, resulting in a conservative sparse network appropriate for the limited sample size (n = 21). The optimal adjacency matrix was used to construct an undirected weighted network using igraph, where nodes represent ASVs and edges represent robust covariance between ASVs. Edges positive and negative weight values indicated respectively co-occurence and co-exclusion relationships. Unconnected nodes were removed before further analysis. Modules (clusters of co-varying taxa) were identified using the Louvain community detection algorithm with cluster_louvain [43], which optimizes network modularity. Network layout was computed using the Fruchterman-Reingold algorithm with absolute edge weights.

To evaluate the specificities of candidate indicator taxa in the network, those taxa were first divided into five categories: associated with faster, or slower, t50germ, or higher, or lower, FEP, or non-indicator. To assess whether taxa from the same association category tend to co-occur, the assortativity of taxa nodes (i.e., the more-than-random occurrence between taxa of the same association category) was computed on each category. Assortativity is a metric bound by [-1, +1], where +1 means that every edge connects two nodes of the same category, 0 that edges connect same-category and different-category nodes at random, and −1 that every edge connects nodes of different categories. Assortativity was computed per category on edge subsets (global: all edges, within-kingdom (bacteria-bacteria and fungi-fungi) or inter-kingdom (bacteria-fungi) edges) using assortativity_nominal from igraph v.2.3.1 [44]. With 21 samples for 189 retained ASVs, network reconstruction is underdetermined and sensitive to sampling variation. Conservative StARS regularization limits spurious edges, but individual edges cannot be interpreted as demonstrated interactions. Modular structure and assortativity patterns are therefore reported as exploratory hypotheses requiring confirmation in a larger sample set.

## Results

### Seed germination and seedling emergence vary between bean varieties

Seed germination time courses were monitored for eight common bean varieties on pleated paper in a germination bench (**Figure 1A-B**). Germination time curves (**Figure 1A**) were used to compute final germination percentage (FGP, **Figure S2A**) and fitted to derive the time to 50% of germinated seeds for each replicate (t50germ) and its variety average (mean t50germ) (**Figure 1B, Figure S2B**). The varieties are separated into two groups. On the one hand, Vezer, Flavert, Deezer, and Facila showed the fast germination, with short lag phases before first germination (22-38 h), lower mean t50germ (32-50 h), and high germination uniformity (17-30h time window between 10% and 90% germination). On the other hand, Vanilla, Caprice, Linex, and Contender showed longer lag phases (37-48 h), higher mean t50germ (54-67 h) and lower uniformity (30-50 h). Mean final germination percentage (mean FGP) also differed between varieties: the four faster-germinating varieties had a mean FGP value ranging from 85.4% to 100%, while amongst slow varieties Contender was notable in reaching a high mean FGP despite its slow germination kinetics.

**Figure 1.**
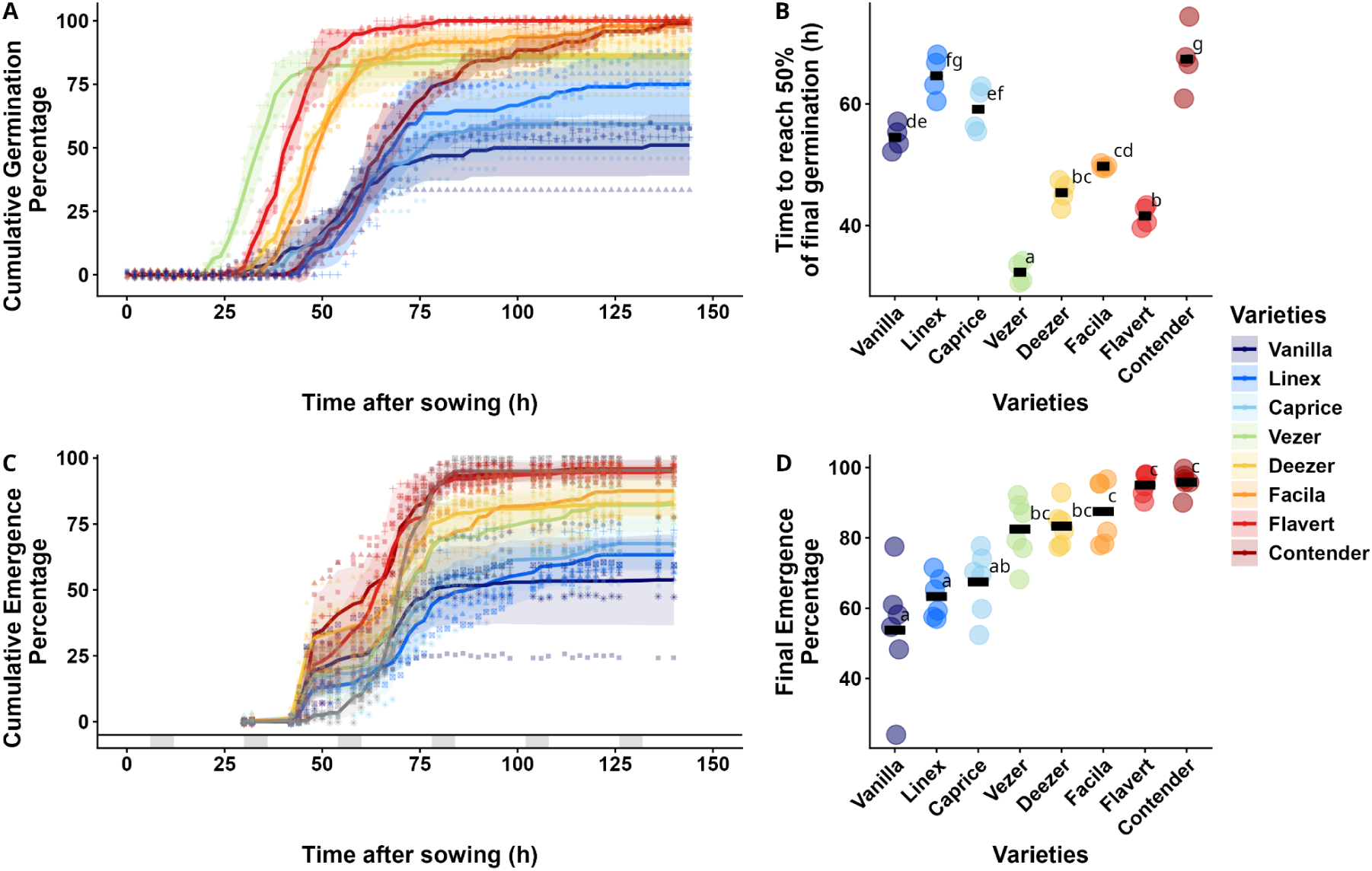
Seed germination and seedling emergence traits of 8 common bean varieties. A: **Cumulated germination percentage over 5 days** (pleated paper on Jacobsen germinators, 20°C, 144 h, 4*24 seeds). Replicates are shown as different shapes, varieties as colors. Bold lines: variety mean; shaded zones: standard deviation interval. B: **Time (h) to 50% germination (t50germ)** estimated by a Four Parameters Hill Function model (data from A, model in **Figure S2B**). Black rectangles: variety mean t50germ. C: **Cumulated emergence percentage over 7 days** (potting mix, 140h, 6*40 seeds). Replicates are shown as different shapes, varieties as colors. Bold lines: variety mean; shaded zones: standard deviation interval. Grey vertical zones above the x-axis indicate night periods. D: **Final emergence percentage (FEP)** at 140 h after sowing (6*40 seeds). Black rectangles: variety mean FEP. Variety colors reflect FEP ranking and are consistent across figures. Different letters indicate significant differences (P < 0.05, one-way ANOVA, post-hoc pairwise comparison of adjusted means).

Seedling emergence was followed during seven days in potting soil (**Figure 1C**). Mean final emergence percentage (mean FEP) differed significantly between varieties, ranging from 54% (Vanilla) to 96% (Contender) (**Figure 1D**). Mean t50germ and mean FGP were not correlated across varieties (**Figure S2C**) whereas mean FGP was highly positively correlated with mean FEP (R^2^ = 0.89, p < 0.001, **Figure S2D**). This suggests that the final germination percentage, but not the germination speed, is a determining factor of seedling emergence of varieties under standard conditions. Based on these results, the mean t50germ and the mean FEP were selected as response variables to which seed metabolic and microbiological characteristics may be associated.

### Between-variety seed size correlates with germination speed while intra-variety seed weight variance correlates with seedling final emergence percentage

Seed weight and size (i.e. projected area) were characterized for individual seeds before imbibition, and their influence on variety seed germination and seedling emergence variability was explored. The eight varieties showed substantial diversity for both traits (**Figure 2, Figure S3A-B**). Variety means ranged from 94 to 522 mg for weight and from 38 to 97 mm² for area. Caprice, Contender and Linex presented the largest and heaviest seeds and the largest intravariety variance for both traits. Conversely, Vanilla and Vezer had the smallest and lightest seeds, with a small variance.

**Figure 2.**
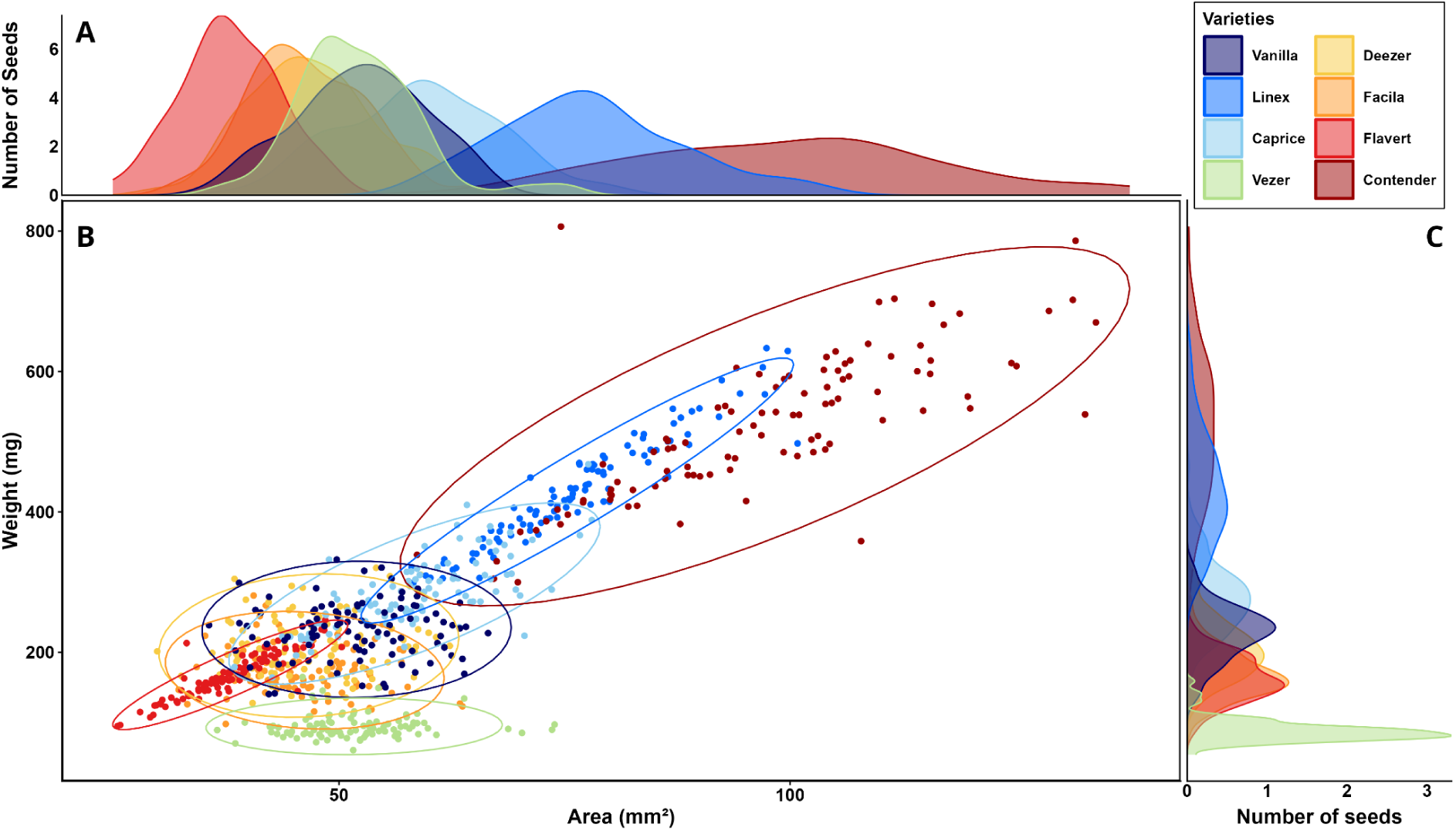
**Weight-area distribution** of individual seeds (100 seeds for each variety) A: **Distribution** of seed area per variety. B: **Weight-area dot plot**. Each dot represents one seed. One color is attributed to each variety according to FEP gradient and is consistent across panels. Ellipses represent 95% confidence intervals based on a multivariate t-distribution. C: **Distribution** of seed weight per variety.

#### Between-variety but not within-variety seed size correlates with t50germ

When varieties were compared (inter-variety analysis), seed weight and projected area were positively correlated with germination speed (R² = 0.94, p < 0.05 and R² = 0.62, p = 0.021 between mean t50germ and mean seed weight or seed area, respectively; **Figure S3D-E**). This indicated that larger and heavier seeds tended to germinate more slowly than small and light seeds. However, Spearman correlations between individual seed weight and germination time within each variety, testing whether heavier or larger individual seeds within a variety germinate faster or slower than lighter or smaller seeds of the same variety, showed a significant positive correlation only in Facila (ρ = +0.37, p = 0.001, padj = 0.0077) and Vanilla (ρ = +0.31, p = 0.019, padj = 0.0755) (**Table S2**). No significant correlation was observed in the other six varieties, even for those presenting the largest seeds (Caprice, Linex, and Contender) which drive the relationship between seed size and mean germination speed when comparing varieties (**Figure S3C**).

#### Intra-variety variance of seed weight correlates with mean final emergence percentage

Unlike germination speed, neither seed weight nor projected area were significantly correlated with emergence (mean FEP) across varieties (**Figure S3F-G**). Given the high disparity in seed weight distribution between varieties, the coefficient of variation of individual seed weight within each variety (referred to as cv_weight) was explored further. Cv_weight did not correlate with mean seed weight itself (ρ = +0.24, p = 0.570), nor with mean germination speed (ρ = +0.21, p = 0.610, **Figure S3H**), but was significantly positively correlated with mean FEP across varieties (ρ = +0.83, p = 0.010, **Figure S3I**, **Table S3**). In other words, varieties with more heterogeneous seed weight distributions, such as Contender and Flavert, tended to achieve higher mean FEP. Conversely, Vanilla, the variety with the most homogeneous seed weights (cv_weight = 0.17), showed the lowest mean FEP (53.75%). Collectively, these results suggest that mean-variety seed weight and size are related to germination speed but unrelated to seedling emergence capacity. Rather, seedling emergence capacity appeared to be linked to intra-variety seed weight heterogeneity.

### Metabolome composition is associated with germination potential

#### Metabolome composition differs across seed organs

Untargeted GC-MS metabolomics was performed on individual seeds dissected into three organs: cotyledon, hypocotyl-radicle axis (HR-axis), and plumule. After quality filtering (removal of samples with null or aberrant internal standard abundance) and normalization by internal standard, 252 annotated metabolites were retained prior to prevalence filtering, for 1 to 12 replicates per organ per variety (**Table S4**, **Figure S4**).

Principal component analysis (PCA) showed a clear difference of clustering in cotyledon versus both plumule and HR-axis. Seed organs were separated along the first two principal components (PC1: 24.0%, PC2: 6.3% of total variance; **Figure 3A**). Organ-level clustering was consistent across varieties, indicating that organ-specific metabolic signature was independent of the genotype. PERMANOVA on Euclidean distances confirmed that organ identity explained 19.4% of total metabolomic variance (F = 16.2, p = 0.001), with this separation concentrated in the major axes of variation (PERMANOVA on PC1-PC2: R² = 0.58, F = 92.2, p = 0.001). The multivariate dispersion differed significantly across organs (betadisper: F = 13.6, p = 0.001), with cotyledon samples being substantially more homogeneous than HR-axis or plumule samples (average distance to group median: 8.98, 13.93, and 14.98 respectively; pairwise tests: Cotyledon vs HR-axis p = 0.001, Cotyledon vs Plumule p = 0.001, HR-axis vs Plumule p = 0.424). That is, there were differences on average between organs along with greater within-organ metabolic variability in the HR axis and plumule, relative to cotyledons.

**Figure 3.**
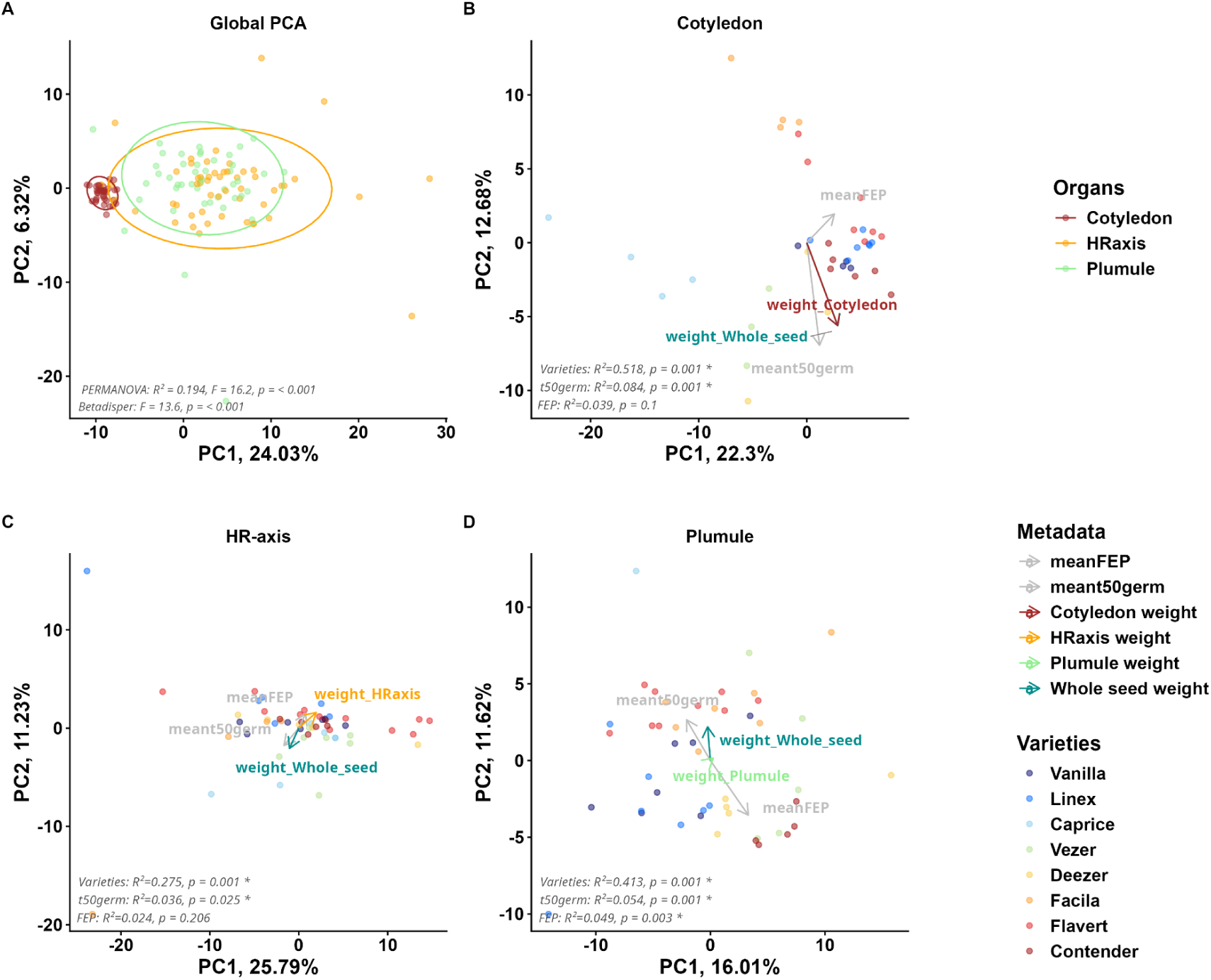
Seed organ metabolome composition. A: **Global principal component analysis (PCA)** of centered-scaled GC-MS metabolite profiles across three seed organs (n = 137 seeds, 221 metabolites after prevalence filtering). Each point represents one individual seed, colored by organ (brown: cotyledon; orange: HR-axis; green: plumule). Ellipses: 95% confidence intervals. PERMANOVA and betadisper statistics are shown in the lower left corner. B-D: **Per-organ PCA** on per-organ Z-scaled metabolome profiles. **B**: cotyledon (n = 38), **C**: HR-axis (n = 53), **D**: plumule (n = 46). Each point represents one individual seed, colored by variety. Arrows indicate the direction and strength of association of continuous variables (seed weight, organ weight, mean t50germ, mean FEP) with the principal component axes, projected with their Pearson correlation with PC1 and PC2 scores scaled by a factor 10 as coordinates. PERMANOVA statistics are shown in the lower left corner.

Metabolites significantly enriched in an organ compared to at least one other organ were identified by lmm tests and Dunn pairwise tests with BH correction. Ten analytes were enriched in **cotyledons**: central carbon metabolites including sugar phosphates (malic acid, ribose-5-phosphate, ribulose-5-phosphate, pyruvic acid, saccharic acid, erythrose-4-phosphate), aromatic compounds (shikimic acid, 2,6-dimethylaniline), and *N*-acetyl-L-alanine (**Table S6**). Six metabolites were enriched in the **HR-axis**, including the allantoin (purine catabolism product), methionine, sinapinic acid, and D-ribose-5-phosphate. Eleven metabolites were enriched in the **plumule**, including compounds of nitrogen metabolism (2,3-diaminopropanoic acid, 4-aminobutanoic acid, citrulline, methionine sulfone), adenine, sucrose, and metabolites of various pathways (heptadecanoic acid, urocanic acid, citramalic acid, maleamate, allocholic acid).

#### Variety drives metabolome composition within each organ

Separate PCA for each organ, colored by variety, showed clear clustering between varieties (**Figure 3B-D**). Within-organ PERMANOVA (**Table S8**) revealed that variety was a main driver of metabolome composition of the three organs, explaining 51.8% of metabolomic variance in cotyledons (F = 4.61, p = 0.001), 41.3% in plumules (F = 3.81, p = 0.001) and 27.5% in the HR-axis (F = 2.44, p = 0.001), with homogeneous dispersion across varieties in all organs (betadisper: Cotyledon F = 1.96, p = 0.094; HR-axis F = 0.91, p = 0.493; Plumule F = 2.07, p = 0.075). Since varieties also differ in their germination and emergence performance, we tested whether these traits were directly associated with metabolome composition, despite the PCA not displaying a clear gradient in mean t50germ or mean FEP along the major axes of variation. PERMANOVA analyses revealed that germination speed (mean t50germ) was significantly associated with metabolome composition of the three organs (mainly in cotyledons: R² = 0.080, F = 3.13, p = 0.001; plumule: R² = 0.056, F = 2.59, p = 0.001 and then HR-axis: R² = 0.039, F = 2.07, p = 0.020). As for mean FEP, it was only significantly associated with the metabolome composition of plumule (R² = 0.067, F = 3.15, p = 0.001) and HR-axis (R² = 0.033, F = 1.75, p = 0.045).

#### Individual seed metabolites associate with t50germ in three organs, and with FEP in two

Candidate metabolites associated with germination speed (t50germ) and emergence (FEP) were investigated through orthogonal partial least squares regression (OPLS) combined with univariate correlation. Model fit was high for t50germ in all three organs (plumule R²Y = 0.956 ± 0.018, Q² = 0.778 ± 0.032; cotyledon R²Y = 0.919 ± 0.055, Q² = 0.699 ± 0.042; hypocotyl-radicle axis R²Y = 0.850 ± 0.025, Q² = 0.575 ± 0.043) and moderate for FEP (Q² = 0.496–0.607; **Table S9**), as evaluated from cross-validation folds drawn at random over individual seeds. However, those high values are due to variety recognition rather than a metabolome-t50germ or -FEP relationship: per-variety t50germ or FEP permutation did not lowered Q² (medians of 0.575-0.766). Under leave-one-variety-out cross-validation, Q² was negative in all six organ x variable combinations (−0.45 to −31.5), fell below the variety-shuffled null in five of six, and was never distinguishable from it (p = 0.19–0.76; **Table S9**). The metabolites below are therefore reported as candidates identified within this panel of eight varieties. From the OPLS, 50, 19 and 12 metabolites were associated with t50germ in cotyledon, plumule and hypocotyl-radicle axis respectively, and 4, 19 and 0 were associated with FEP (**Table S10**). Loading direction and significance were stable across permutations for all retained candidates (CV of pq1 0.03–0.19; CV of −log₁₀ p 0.07–0.31). Of these, 12, 13, and 7 candidate biomarkers for t50germ, and 2 and 14 for FEP, were also supported by at least one independent approach (**Figure 4**).

**Figure 4.**
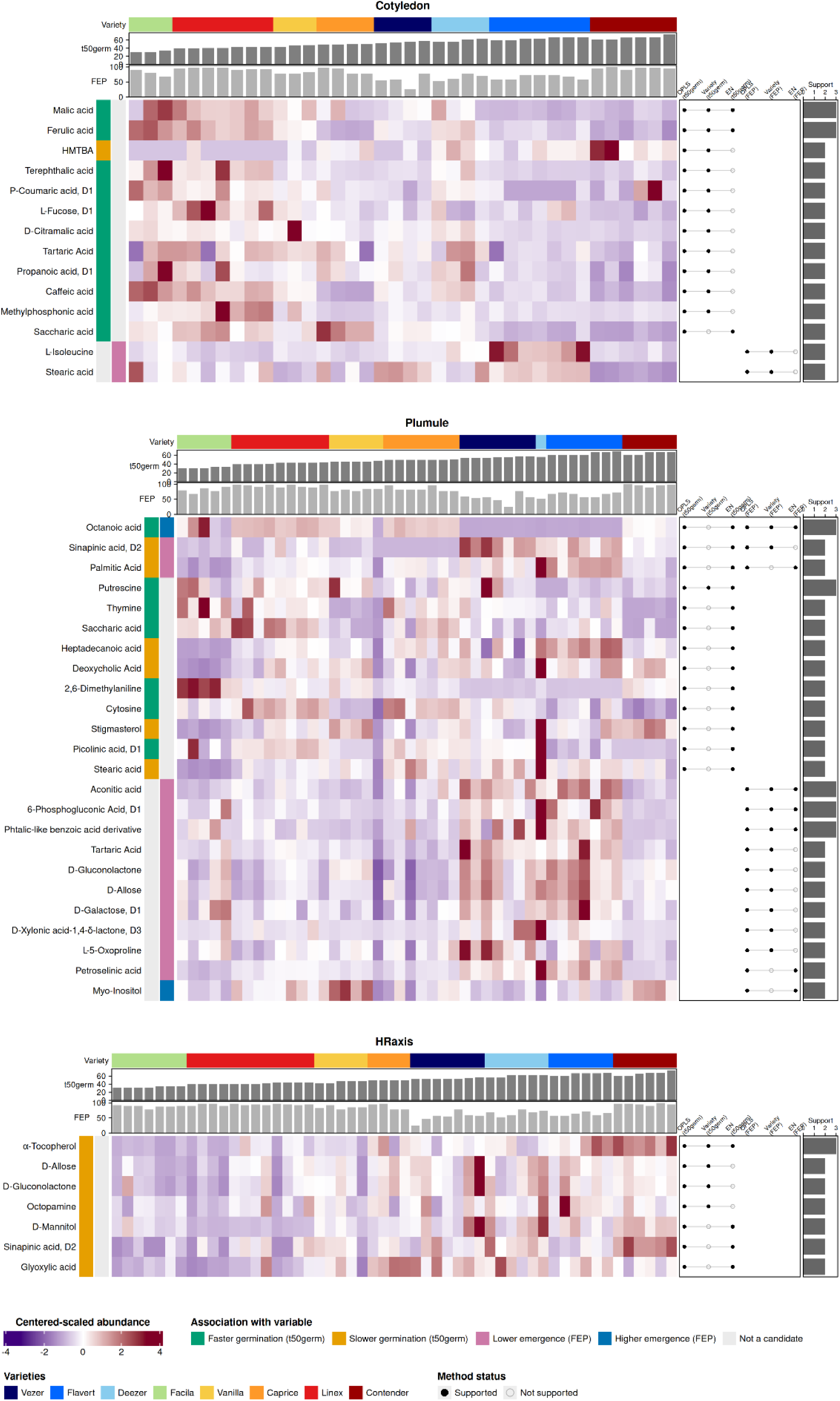
**Heatmap of the candidate metabolite biomarkers of germination speed (mean t50germ) or emergence percentage (mean FEP)** identified by the OPLS permutation pipeline, filtered by stability criterion, and supported by at least two approaches: in the cotyledon (top, 12 metabolites for t50germ, 2 for FEP), plumule (middle, 13 metabolites for t50germ, 14 for FEP), and hypocotyl-radicle axis (HRaxis, bottom, 7 metabolites for t50germ). Rows: metabolites (hierarchical clustering, Euclidean distance); columns: individual seeds ordered by variety (color bar, top) then by individual t50germ within variety (grey barplot, top; higher bars = slower germination). Metabolite abundance is shown as per-organ centered-scaled values (purple: below organ mean; white: at organ mean; red: above organ mean). Left strips: sign of association with t50germ (green: faster germination; orange: slower germination) or FEP (pink: lower FEP, blue: higher FEP). Right panel: approaches supporting each metabolite (OPLS permutation frequency, variety-level Spearman correlation, and elastic net regression); filled circles: support. Barplot: maximum number of confirming methods (2-3). Candidates are hypothesis-generating: leave-one-variety-out cross-validation and a family-level permutation test did not support generalisation beyond the eight varieties.

In cotyledons, eleven out of the twelve candidate biomarkers associated with t50germ were more abundant in faster-germinating varieties. They comprise organic acids of central carbon metabolism (malic acid (ρ = −0.93, p = 0.0009), citramalic, tartaric and saccharic acids), hydroxycinnamic acids of the phenylpropanoid pathway (ferulic acid (ρ = −0.77), p-coumaric (ρ = −0.82) and caffeic acids) together with L-fucose, terephthalic, propanoic and methylphosphonic acids. The single opposite-direction candidate was HMTBA (2-hydroxy-4-methylthiobutyric acid, ρ = +0.85), a methionine analogue, more abundant in slower-germinating varieties. Cotyledon also harboured two candidate biomarkers associated with lower FEP: L-isoleucine (ρ = −0.90, p = 0.002) and stearic acid (ρ = −0.79). In plumule, putrescine was the only candidate biomarker for t50germ supported at the variety level (ρ = −0.89, p = 0.003), and was more abundant in faster-germinating varieties.

Among the twelve other candidate biomarkers, saturated fatty acids (palmitic, stearic, heptadecanoic) and sterols (stigmasterol, deoxycholic acid) were more abundant in slower varieties, pyrimidine bases (thymine, cytosine) and short-chain acids in faster-germinating varieties. Plumule also harboured fourteen candidate biomarkers associated with FEP. Twelve were more abundant in varieties with lower FEP: sugars and sugar derivatives (D-allose, D-galactose, D-gluconolactone, D-xylonic acid-1,4-δ-lactone), the oxidative pentose-phosphate intermediate 6-phosphogluconic acid (ρ = −0.81), organic acids (tartaric, ρ = −0.90; aconitic, ρ = −0.82), sinapinic acid, L-5-oxoproline and two fatty acids (palmitic, petroselinic). Only two candidate biomarkers were more abundant in high-FEP varieties: octanoic acid (ρ = +0.90, p = 0.003) and myo-inositol. In hypocotyl-radicle axis, the seven candidate biomarkers were more abundant in slower-germinating varieties: α-tocopherol (ρ = +0.88, p = 0.004), D-allose, D-gluconolactone, octopamine, D-mannitol, sinapinic acid and glyoxylic acid.

### Seed microbiota is associated with variety’s identity and final emergence percentage

#### Seed microbiota is dominated by a few bacterial and fungal genera

Illumina sequencing yielded an average of 45,050 raw reads of *gyrB* gene sequences and 29,445 of ITS gene sequences, respectively. The sequences were filtered and merged to an average of 11,408 final bacterial and 16,870 final fungal reads per sample, from which 4,789 bacterial and 357 fungal ASVs were identified. Reads were rarefied to 8,975 reads per sample (99.98% coverage) for gyrB and to an even depth of 7,625 reads per sample for ITS, resulting in 4,789 bacterial and 346 fungal ASVs retained after rarefaction ( **Figure S5.G-H**). The most prevalent genera were *Pantoea*, *Pseudomonas*, *Kosakonia*, and *Erwinia*, comprising in average more than 54% of the relative abundance in the bacterial communities, and *Alternaria*, *Cladosporium*, and *Stemphylium*, comprising in average more than 90% of the relative abundance in the fungal communities (**Figure S5.A-D**).

#### Fungal, but not bacterial, community composition is variety-dependent

α-diversity varied significantly between varieties from the tested seed batch for both bacterial and fungal communities (Kruskal-Wallis, p = 0.034 and p = 0.025, respectively). β-diversity analysis revealed contrasting patterns (**Figure 5A-B**): variety had no significant effect on bacterial community composition (PERMANOVA: R² = 0.37, p = 0.4) but strongly structured fungal communities (R² = 0.80, p = 0.001), though the low number of replicates per variety (n = 2–3) prevented identification of individually significant variety pairs. For further details on general microbiota composition, see **Note S1**.

**Figure 5.**
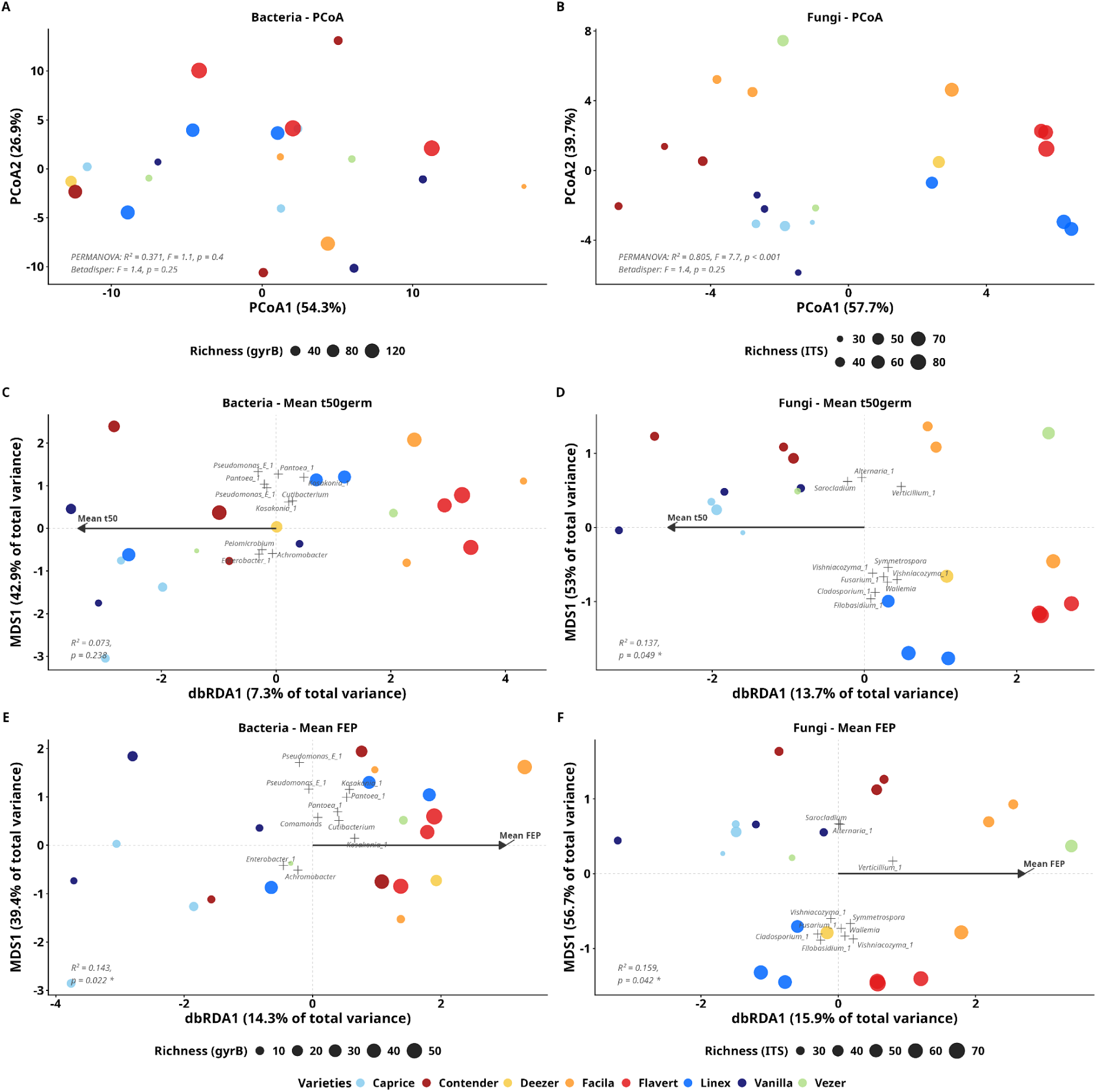
Native seed microbiota community composition and association with germination and emergence traits. A-B: **Principal Coordinates Analysis of bacterial** (A) **or fungal** (B) **community profiles** (Aitchison distance, rCLR transformation). Points: individual samples (n = 21) colored by variety; size: ASV richness. C-F: **Distance-based redundancy analysis (db-RDA)** of bacterial (C, E) and fungal (D, F) communities constrained by mean germination time (mean t50germ) (C-D) or mean final emergence percentage (mean FEP) (E-F). Arrows: constraint direction and strength. Axis labels: percentage of total inertia explained. Grey axis labels: non-significant axes. Species scores were projected post-hoc onto the ordination space.

#### Fungal communities associate with both t50germ and FEP, and bacteria only to FEP

Seed bacterial community composition (*gyrB*) was not significantly associated with mean t50germ (PERMANOVA: F = 1.50, R² = 0.073, p = 0.238) but was associated with mean FEP (PERMANOVA: F = 3.18, R² = 0.143, p = 0.022). In contrast, fungal community composition (ITS) was significantly associated with both traits, although marginally (PERMANOVA: F = 3.01, R² = 0.137, p = 0.049; and F = 3.58, R² = 0.159, p = 0.042, respectively). Because variety identity accounted for the largest share of fungal community variation (R² = 0.80), trait-related compositional differences cannot be statistically disentangled from the variety background. db-RDA biplots were used to visualize these associations, with the ten ASVs most strongly driving the constrained axes identified and labeled by their lowest available taxonomic assignment (**Figure 5C-F, Table S11**).

These results suggest that seed-associated fungal communities are more broadly linked to t50germ and FEP than bacterial communities, which appear specifically associated to FEP rather than to t50germ.

#### Individual fungal taxa are associated with FEP rather than t50germ, and bacteria with both

Individual microbial taxa associated with germination and emergence traits were identified in an exploratory analysis with five independent statistical methods (Spearman rank correlation on replicates or on variety means, linear model-based association analysis, compositional correlation analysis, and bias-corrected compositional differential abundance analysis). Across the five statistical methods applied, a total of 23 taxa reached the exploratory significance threshold (q ≤ 0.25) (**Table S12**). Of these, seven bacterial taxa and four fungal taxa were confirmed by at least two independent methods, and retained as robust candidates (**Figure 6**). For fungal communities (ITS1), a *Stemphylium trifolii*, one unassigned fungi and an unassigned *Saccharomycetales* ASVs showed strong positive associations with FEP (ρ = 0.63, 0.62 and 0.62), one *Acremonium* ASV showed a negative association with FEP (ρ = −0.60). No fungal ASV was detected as robustly associated with low t50germ. For bacterial communities (*gyrB*), an unassignable *Actinomycetes* ASV negatively associated with mean t50germ (ρ = −0.63, confirmed by Spearman, Maaslin and ALDEx2), suggesting faster germination in seeds harbouring this taxon. *Kosakonia cowanii*, one *Erwinia* ASV, and a *Reyranella soli* were positively associated with mean FEP, as confirmed by Spearman correlation and ANCOM-BC2 or mean Spearman correlation (ρ = 0.70, 0.56, and 0.58 respectively). As such, seeds harbouring higher abundances of these bacteria tended to germinate faster.

**Figure 6.**
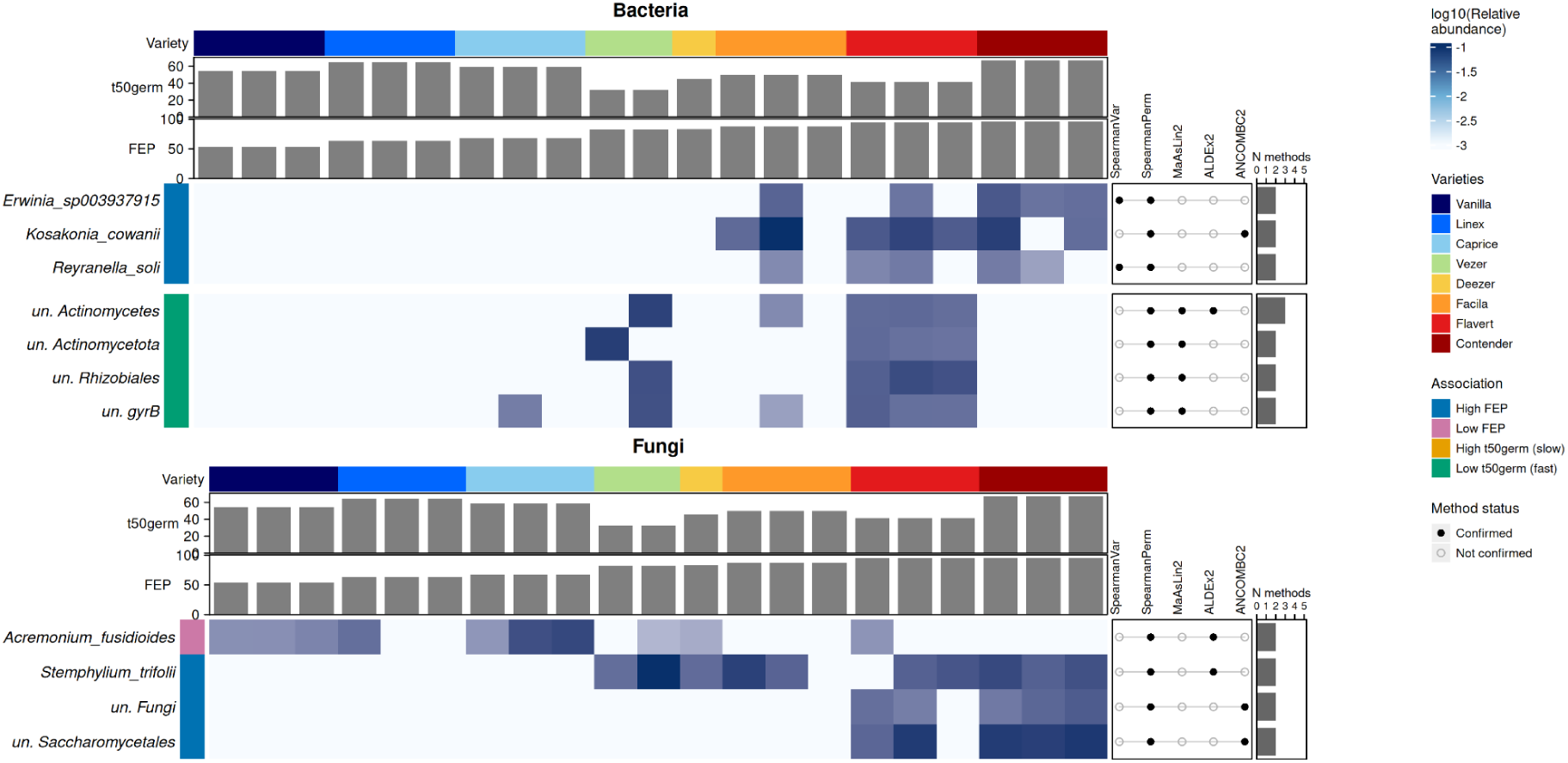
Heatmap of the 13 taxa associated with germination speed (t50germ) or emergence percentage (FEP) identified by at least 2 methods. Rows: taxa; columns: individual seed batch microbiota ordered by variety (color bar, top) then by replicate FEP within variety (lower barplot). Microbial taxa abundance TSS-transformed is shown on a log_10_ scale. Left color strip: sign of association with t50germ or FEP (green: faster germination; blue: lower FEP, pink: higher FEP). Right panel: confirmation across five methods (Spearman correlation (rho) on variety-mean values and on per-replicate permutations, MaAsLin2 (log fold-change), ALDEx2 (rho), and ANCOM-BC2 (log fold-change)); filled circles: confirmed. Barplot: number of confirming methods. All associations are exploratory (q ≤ 0.25).

Notably, *Kosakonia cowanii* was identified both as a robust indicator taxa associated with mean FEP from two methods and as one of the 10 taxa most strongly driving the db-RDA axis associated with mean t50germ and mean FEP. A one-method candidate for FEP *Cladosporium herbarum* (ALDEx2 only, ρ = −0.69 negative) also appears to drive both fungal dbRDA ordination.

#### Fungi with similar associations to t50germ or FEP preferentially co-occur

To evaluate the tendency of taxa to connect preferentially to other taxa with similar association with t50germ or FEP, a multi-kingdom co-occurrence network was inferred (stability = 0.042, optimal lambda = 0.606). After removal of isolated nodes, 184 taxa remained in the connected network with 247 edges (101 bacteria, 83 fungi, **Figure 7A**). Edge weights ranged from −0.278 to 0.422, with positive weights indicating positive covariance (163 edges) and negative weights indicating exclusion between taxa (84 edges) (**Table S13**).

**Figure 7.**
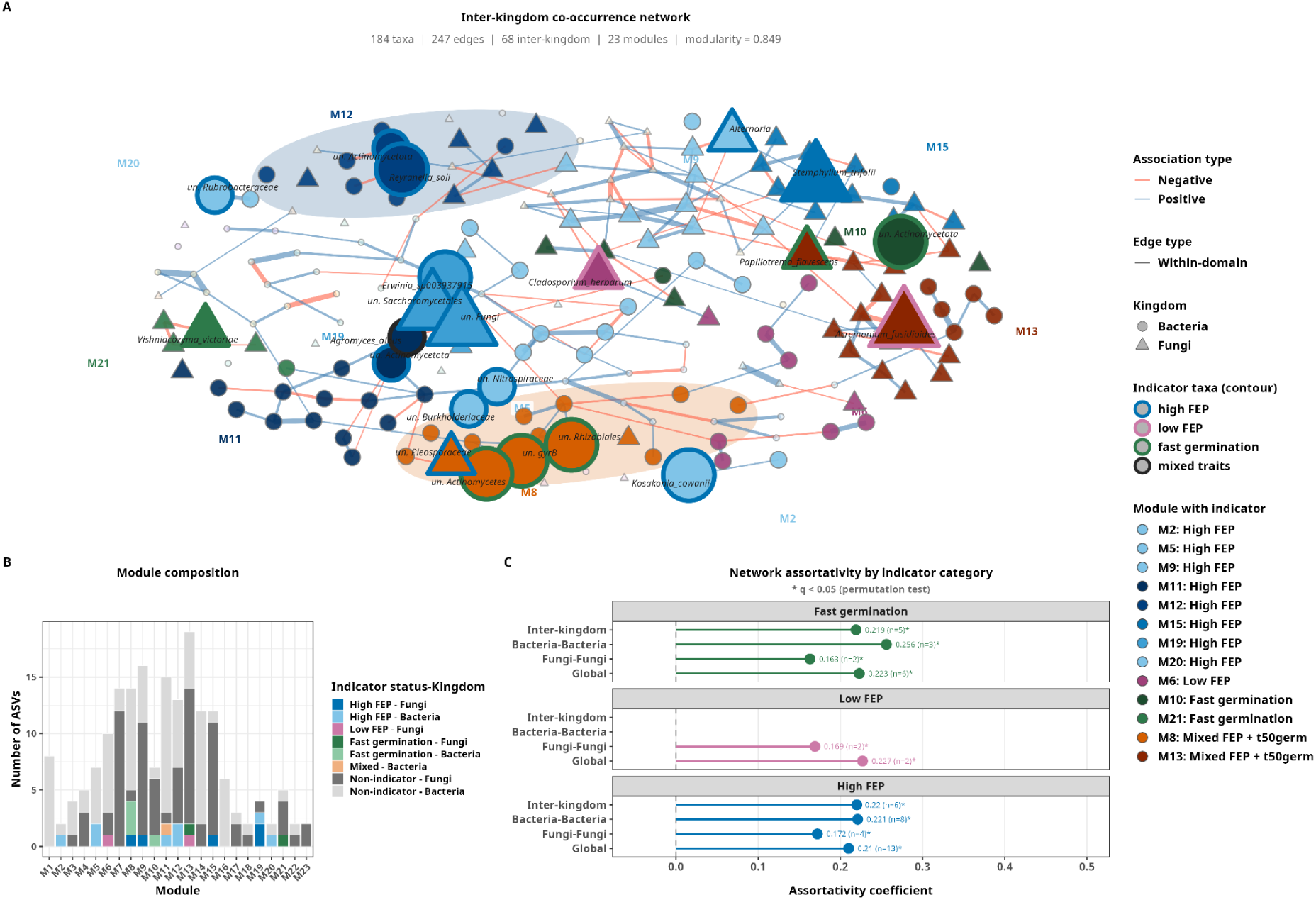
Seed microbiota co-occurrence network and module composition. A: **Inter-kingdom co-occurrence network** inferred by cross-domain SPIEC-EASI (ASVs with prevalence > 15%). Nodes: ASVs; size: indicator status (large with contour: candidate indicator taxa identified by several (biggest) or one (medium) method, small: taxa in modules containing indicator taxa, tiny grey: others). Candidate indicator taxa contours are colored by association (high FEP: blue; low FEP: pink; high t50germ - slow germination: orange; low t50germ - fast germination: green). Nodes fill color: Louvain module (bright colors: modules containing indicator taxa: high FEP: blue shades; low FEP: pink; high t50germ: yellow; low t50germ: green; FEP and t50germ mixed: orange/red shades; high and low t50germ: cyan; high and low FEP: purple). Edge color and style: sign of the partial correlation between taxa (blue: positive, red: negative, solid: within-domain, dotted: inter-kingdom). Isolated nodes are not shown. *un. : unassigned*. B: **Network module composition** by kingdom, evidencing candidate indicator taxa associated with each category (high or low t50germ, or high or low FEP). C: **Network assortativity** with assortativity coefficient for each trait (All traits, t50germ, FEP) computed across different edge subset (Global: all edges; Bacteria-Bacteria and Fungi-Fungi: within-kingdom edges; Inter-kingdom: edges between bacterial and fungal taxa) for each association sign. Positive values: taxa from the same association category tend to co-occur. Absence of value corresponds to non-occurring associations (e.g., for slow-germination-associated taxa). Node and edge counts per subset are indicated in grey.

Taxa correlating with FEP or t50germ were distributed across multiple modules. Fungal indicator taxa of the same association category (associated with faster or slower t50germ, or higher or lower FEP) appeared in several modules (including dark blue M12 on top of **Figure 7A**, **Figure 7B, Table S13**). The mixed modules (orange/red : containing both mean t50germ and mean FEP-related taxa) suggest that some community members had a role in both germination and emergence traits. Note that one candidate taxa associated with higher FEP, *Kosakonia cowanii*, was removed from the network as it was not connected to any other taxa.

Assortativity coefficients (**Figure 7C**) revealed that indicator taxa co-occured with other indicator taxa of the same category more than expected by chance for three of five categories tested. This was strongest for high-FEP-associated taxa (global assortativity = 0.21, z = 4.58, p = 0.001), and also significant for low-FEP-associated taxa (global assortativity = 0.227, z = 4.11, p = 0.001) though based on fewer nodes. For both high-FEP and fast-germination categories, assortativity was significant within each kingdom separately (bacteria-bacteria: 0.221 and 0.256 respectively, fungi-fungi: 0.172 and 0.163) as well as across-kingdom (0.22 and 0.291), with all q-values ≤ 0.03 after Benjamini-Hochberg correction. No significant same-category co-occurrence could be tested for slow-germination associated taxa (high t50germ), as none of these indicator taxa were retained as connected nodes in the network, nor for taxa with mixed trait associations, which comprised too few nodes (≤ 1 per edge subset) to compute assortativity.

## Discussion

### Germination speed and seedling emergence capacity are independent in common bean

The distinction between germination and emergence is well established, with extensive experimental and conceptual work showing that these two phases rely on partly distinct underlying processes and determinants in crop and model species [45–48], and that rapid germination does not guarantee successful seedling establishment [49, 50]. Contender, the slowest-germinating variety achieving the highest emergence percentage (95.8%), illustrates this functional decoupling and highlights the risk of using germination speed as the sole proxy for seed quality in field contexts. Building on this framework, we investigated quantitative candidate indicators in common bean associated with early germination processes (metabolic activation, radicle protrusion) on the one hand, and traits associated with seedling emergence (hypocotyl elongation, ability to overcome soil resistance, and early seedling establishment) on the other hand. The aim of this study was to identify practical, physiologically grounded candidate indicators of seed quality, using separately seed germination and seedling emergence as proxies, emphasizing the importance of both processes in seed testing rather than relying on a single measure of seed quality that aggregates germination and emergence. To do so, we used accurate, individual-seed weight and morphological measurement, a fine individual-seed per-organ metabolome characterisation, and seed batch microbiota analysis for eight varieties of common bean (*Phaseolus vulgaris* L.). We took the precaution to study seeds produced in the same assay so that the seed microbiota composition and seed quality would reflect varieties-related properties rather than environmental variations between location, and timing of seed production. The eight varieties, representative of the morphological and genetic diversity of cultivated varieties in France, display a wide range of germination speeds and emergence capacities we used to identify proxies of seed quality. Given the agronomical importance of such proxies, their strength and reliability should be validated on seeds of a larger panel of varieties produced in different locations, and further evaluated across various conditions in germination and emergence assays. Indeed, the growth media and conditions used in germination and emergence assays can impact germination timing, as shown with Contender, that already reaches 50% of emergence in potting soil at 78 h, while 50% of germination in pleated paper at only 76 h.

### Seed size correlates with germination speed between varieties, while intra-variety size variance reflects emergence

When varieties were compared, we found that seed weight and size were good predictors of germination speed (e.g. weight vs mean t50germ: R² = 0.94), which is consistent with reports in other legume species where large seed mass leads to slower water uptake kinetics and longer imbibition times [51–54]. We nevertheless recognise that it is not a simple correlation and some legume species are associated with an opposite, or more complex, relationship [47]. Also, the relationship between seed weight and t50germ was not found anymore when individual seeds, rather than variety averages, were considered. Thus, it seems that in common bean, the mass-germination time relationship was a statistical property at the variety scale, rather than a reflection of intrinsic mechanical effects of seed mass on, e.g., imbibition. In other words, differentiating varieties encompasses not only morphological differences (such as average seed weight), but also physiological differences that dictate germination timing in this species. Quite surprisingly, a positive correlation was found between intra-variety seed weight variation and mean FEP (ρ = +0.83, p = 0.010), independent of mean seed weight itself (ρ = +0.24, p = 0.570) (**Figure S2I**). From a physiological perspective, it might reflect a decoupling during seed development, between reserve accumulation and cotyledon development on the one hand, and the biosynthesis of cellular actors involved in germination processes (such as rRNA accumulation, hormone content, etc.) on the other hand. That is, varieties associated with high FEP might give priority to the development of the molecular machinery for germination, regardless of natural seed biomass variations, due to, e.g., variable phloem transport rate, pod position on the plant, or seed position within the pod [28]. From an ecological point of view, it could suggest that varieties producing more heterogeneous seed populations are better adapted to unpredictable soil conditions during emergence. This would be in line with the concept of “bet-hedging” in seed biology, whereby variability in morphology and developmental stage within a seed population ensures a higher success rate of emergence under variable environmental conditions [55, 56].

### Metabolomes of seed organs provide candidate indicators of germination speed and FEP

As for proteins and transcripts, metabolites accumulated during seed maturation and preserved during desiccation can play a role in germination [57]. Seed organs have different physiological functions, reflected in their composition [58–60]. Here, the three dissected seed organs exhibited distinct metabolomic profiles that were consistent with their developmental functions. The cotyledons were enriched in metabolites of primary carbon metabolism including sugar phosphates (reflecting sugar interconversions and starch accumulation), as well as pyruvate and malate (glycolysis and Krebs cycle) and *N*-acetyl-alanine (stabilisation of accumulated proteins) [61]. The hypocotyl-radicle axis was enriched in allantoin, which is a nitrogen transport metabolite in French bean [62], and methionine, a sulphur transport molecule re-synthetised from S-methylmethionine in legume developing seeds [63]. The plumule was enriched in amino acids, adenine, polyamine (citrulline) and sucrose, reflecting metabolism sustaining leaf cell division and growth [13, 64].

Within each organ, variety was a dominant driver of the metabolome (R² = 0.28-0.52), confirming that seed metabolomic profiles were primarily dictated by the genotype, as found previously in common bean [4], in tomato [7, 65] but not in *Medicago* [28]. Candidate biomarkers were identified for both t50germ and FEP, and differed in their organ distribution and chemistry. Germination speed yielded candidates in all three organs, whereas emergence capacity yielded them only in the plumule and cotyledon, and none in the hypocotyl-radicle axis. This contrast, together with the near absence of overlap between the two candidate sets, is consistent with germination speed and emergence capacity relying on distinct metabolic bases. In **cotyledons**, the storage organ mobilized during imbibition, candidate biomarkers of faster germination were dominated by organic acids of the central carbon metabolism, and by hydroxycinnamic acids. Malate is a central organic-acid intermediate connecting mitochondrial respiration, anaplerotic metabolism and, in oil-rich tissues, glyoxylate-cycle-associated reserve mobilization during early heterotrophic growth. Its greater abundance in faster-germinating varieties is therefore consistent with a metabolite state compatible with rapid respiratory activation at imbibition. D-citramalate is formed by condensation of acetyl-CoA with pyruvate and is converted, via citraconate, to 2-oxobutanoate, a precursor of isoleucine. Its accumulation therefore reports on the availability of the pyruvate and acetyl-CoA pools that supply respiration at imbibition. Ferulic and p-coumaric acids can occur as ester-linked hydroxycinnamate modifications of plant cell-wall polymers, where oxidative coupling may increase wall cross-linking and reduce extensibility [66]. In dicots, ferulate has also been implicated in pectic and extensin-associated cross-links [67]. Phenolic composition and wall mechanics can plausibly influence hydration and embryo expansion. In the **hypocotyl-radicle axis**, the organ whose elongation drives radicle protrusion, all seven candidate biomarkers were more abundant in slower-germinating varieties and formed a coherent antioxidant and osmolyte signature. α-tocopherol is the principal lipid-soluble antioxidant of seed tissues. Its increase during early imbibition has been observed alongside ROS accumulation and lipid peroxidation, consistent with recruitment of antioxidant protection [68, 69]. Gluconolactone, the product of glucose-6-phosphate oxidation by glucose-6-phosphate dehydrogenase, marks flux through the oxidative branch of the pentose phosphate pathway, the principal source of the NADPH that sustains the glutathione–ascorbate cycle and hence the regeneration of α-tocopherol. Mannitol can act as an osmoprotective compatible solute and has been implicated in oxidative-stress protection, while alloseis a rare aldose whose accumulation has been associated with perturbed hexose metabolism. Taken together, these candidates point to a coordinated investment in oxidative protection and in the reducing power that supports it. A greater constitutive investment in oxidative protection in the axis of slower varieties may reflect a higher basal oxidative burden, a recognised constraint on the speed of germination [70, 71]. Interestingly, putrescine was identified as the most robust plumule candidate biomarker of fast germination regardless of the statistical method used. It has been shown that putrescine plays a role in stress mitigation [72], seed germination [73–76] and growth of young soybean seedlings [77], and its accumulation in the embryonic shoot of rapidly germinating varieties is consistent with an axis primed for cell division and elongation. Thymine and cytosine were also more abundant in faster-germinating varieties, and probably reflect cell division and thus nucleotide synthesis capability. Conversely, free saturated fatty acids (heptadecanoic, palmitic), and stearic acids were more abundant in plumules of slower-germinating seeds, perhaps reflecting increased cell membrane rigidity, improper fatty acid composition balancing, or altered carbon partitioning towards lipid at the expense of carbohydrates and proteins. A link with membrane properties would be consistent with the documented role of membrane fluidity in controlling turgor-driven cell expansion required for radicle emergence [78] and the key role of changes in fatty acid composition during imbibition, as found in *Medicago* [79].

Candidate biomarkers of emergence capacity were chemically distinct from those of germination speed and were concentrated in the plumule. Twelve of the fourteen plumule candidates were more abundant in varieties with lower emergence and comprised free sugars and sugar derivatives, the oxidative pentose-phosphate intermediate 6-phosphogluconic acid, organic acids and L-5-oxoproline, an intermediate of glutathione turnover. The combined accumulation of free reducing sugars, sugar acids and markers of oxidative and glutathione metabolism is frequently observed during seed ageing and loss of vigour. Its concentration in the plumule rather than in the storage tissue may suggest that the embryonic shoot, the organ that must elongate through the soil column, could be one site where emergence capacity would be compromised. In cotyledons, the two supported candidates of lower emergence were L-isoleucine and stearic acid, and the two further candidates supported by the OPLS alone were the branched-chain amino acids leucine and valine, suggesting that free branched-chain amino acid accumulation in the storage tissue may also accompany reduced emergence.

With eight varieties and more than two hundred metabolites, individual associations of the observed strength arise readily by chance, and the argument for the candidates listed here rests on their directional consistency within an organ and on their membership of related pathways rather than on individual statistical support. The stability of such candidate indicators should be verified across a broader diversity of varieties but also across seed size diversity, as the seed metabolome is impacted by seed weight [27]. In this exploratory study, only seeds of weight close to mean weight per variety (± 50% standard deviation around the mean seed weight) were analyzed to account for such weight impact, potentially masking some candidate indicators.

### Fungal community composition is broadly linked to germination performance while bacteria community is more specifically associated with final emergence percentage

Seed microbiota role in germination and seedling emergence is a recent field: a growing number of studies focus on seed microbiota characterisation, while few link that diversity to plant development [80–82]. In the present study, fungal community composition was variety-dependent for the studied seed batches (PERMANOVA R² = 0.80), whereas bacterial community composition showed no significant variety effect (R² = 0.37, p = 0.4). The higher impact of host variety, or species, on fungal communities has already been described [83–85]. This contrast may reflect differences in transmission mode and community assembly, although these mechanisms are yet poorly understood. The dominance of *Pantoea*, *Pseudomonas*, *Kosakonia* and *Erwinia* in the bacterial communities, and of *Alternaria*, *Cladosporium* and *Stemphylium* in the fungal communities, is consistent with the core seed microbiota composition reported across crop species in a meta-analysis [85].

The composition of the fungal community was significantly associated with both t50germ and FEP (PERMANOVA R² = 0.137 and 0.159), while bacterial communities were associated with FEP only (R² = 0.143, p = 0.022). While those exploratory associations need further replication and experimental validation, and cannot be discriminated from variety effect, the dissociation between t50germ and FEP associations could suggest that fungi may be more closely associated to the biochemical activation of germination, possibly through interactions with seed coat chemistry or plumule metabolism, while bacteria may influence the later stages of seedling emergence and establishment, consistent with their documented roles in phytohormone production, phosphate solubilisation and induced systemic resistance [86–88]. Study of the seed coat metabolome, that we were not able to analyse in this study, could provide insight into a potential fungal-tegumen interaction. The positive association of *Saccharomycetales* ASVs with final emergence (ρ = +0.62) could incite further research on the growth-promoting potential of seed yeasts [89, 90]. One *Kosakonia cowanii* ASV was found positively associated with emergence (ρ = +0.70, two independent methods) and identified as a top driver of the db-RDA community axes, providing evidence from community-level and taxon-level analyses. This is coherent with the plant-growth promoting potential reported for this species [91]. Similarly, a *Cladosporium herbarum* ASV, associated with low FEP (ρ= −0.69, 1 method), was in the top db-RDA taxa. This species has been shown to negatively impact *Phaseolus vulgaris* development [92, 93]. Those exploratory associations and the mechanism of their potential impacts on germination and emergence should be further investigated, for example through seed inoculation with isolated microbial strains [94].

Co-occurrence networks offer a way of characterising patterns of taxon association across samples, as well as generating hypotheses about the ecological processes that structure microbial communities. Although they do not reflect direct, biotic interaction, the edges that link taxa nodes allude to facilitation, mutualism, and niche sharing for positive co-occurrence or to the indirect consequences of competition, predation, and environmental filters for negative co-occurrence [95–98]. In our case, the cross-domain co-occurrence network revealed that the seed microbiota is structured with respect to germination performance: fungi with similar exploratory associations with FEP preferentially co-occur. Such association of several candidate indicator taxa in the same modules weights their value as individual indicator, but also of the whole module as a stronger proxy to evaluate the seeds emergence capacities. Indeed, identification of microbial modules associated with ecosystemic functions is a promising research avenue to both find indicators and better understand microbiota contribution to those functions [99–101]. Such modules’ role in seedling emergence can then be validated with inoculation approaches, using microbial consortia (also termed “synthetic communities”) inoculation on seeds to infer causal relationships in plant-microbiota interactions [102, 103].

## Conclusion

Taken together, those results across eight common bean varieties demonstrate that germination speed and emergence capacity in common bean are associated with distinct and largely independent seed properties, each with specific morphological, metabolic and microbial signatures. Seed size and plumule metabolome composition were informative candidate indicators of germination speed, while intravariety seed weight heterogeneity and fungal and bacterial community composition were found informative for emergence. This multi-factor picture supports the development of complementary rather than single-proxy seed quality assessment frameworks, potentially integrating morphometric, metabolomic and microbiota measurements on dry seeds prior to sowing [104, 105]. Future research involving larger variety panels, seeds of diverse production location, individual seed phenotyping designs, and spermosphere analysis at the seed-to-soil interface [106, 107] will be needed to validate the candidate indicators and assess their applicability across environments and agronomic contexts.

## Declarations

### Ethics approval and consent to participate

Not applicable

### Consent for publication

Not applicable

### Availability of data and materials

The datasets generated and analyzed during current study are included as Supplementary material, or can be found with codes used for this manuscript analysis, on GitHub at this address:

https://github.com/LounaCS/Multidimensional-Drivers-of-Seed-Germination-and-Emergence.git

### Author’s contributions

C.-E. K. directed the seed harvest. B.T., E.P., M.S., G.T., and M.B. designed the experiments. C.M., A.D., and M.-A.W. performed the germination assay. L.L.C., and S.H. performed the X-ray analyses. L.C.-S. performed the emergence assay. E.P., C.A. dissected the dry seeds and prepared the metabolomic experiment, C.A. and J.L. ran the metabolomic experiment, and L.C.-S. processed the raw metabolomic data, with guidance from C.A. and G.T.. M.S. and C.M. performed the metabarcoding assay. L.C.-S. analyzed all experimental data. L.C.-S, G.T., B.T., E.P. and M.S. were involved in the interpretation of the results. L.C.-S wrote and revised the article with B.T., E.P., G.T., and M.S. All authors read, corrected, and approved the final manuscript.

## Acknowledgements

This research was conducted in the frame of the 3rd Programme for Future Investments (France 2030), operated by the SUCSEED project (ANR- 20- PCPA-0009) and funded by the ‘Growing and Protecting crops Differently’ French Priority Research Program (PPR-CPA), part of the national investment plan operated by the French National Research Agency (ANR).

We thank Elise Vendeuvre for the gift of Linex, Deezer, Vezer, and Caprice commercial seeds used as mother-plants. We thank Fernand Roques, Elodie Gauvin, and Emmanuelle Laurent for the seeds they grew and provided, and Jérôme Lemaire for cleaning and sampling seed lots.

We thank Didier Demilly and the Phenotic Angers Seed & Plant Phenotyping Facility (INRAE-IRHS, Angers University, Institut Agro, GEVES) for experiments on seed germination and seedling emergence.

We thank Pascale Satour for her contribution to seed dissection and metabolite extraction, Anne Préveaux for her contribution to sample processing for microbiota analysis, and Aurélie Charrier for her contribution to the X-ray analyses.

L.C.-S would like to thank Agathe Brault, Chrystelle Brin, Louis Broussard, Thomas Chadelaud, Armelle Darrasse, Isa Hollop, Bastien Gouffier, Marie-Agnès Jacques, Oscar Joubert, Logan Suteau, Boris Taillefer, and Ismail Zaag for their help during the emergence assay, and Kenji Maurice for the discussions on networks.

## Supplementary material

**Supplementary Note**: pdf file with methods and results on microbial community composition, evaluated with classical α- and β-diversity metrics.

**Supplementary Figures**: pdf file with complementary figures:

Figure S1: *Supplementary data on experimental design,*

Figure S2: *Supplementary data on Germination and Emergence*,

Figure S3: *Supplementary data on Morphological characteristics*,

Figure S4: *Heatmap of the relative metabolic abundance of the samples*,

Figure S5: *Supplementary data on seed microbiota characteristics*.

**Supplementary Tables**: excel file summarizing the datasets and statistical results for each analyse.

Supplementary Table 1: *Morphological mean values per variety*

Supplementary Table 2: *Within-variety Spearman correlations between morphological variables and imbibition time at the single-seed level*

Supplementary Table 3: *Between-variety correlation between morphological variables and meanFEP or meant50germ*

Supplementary Table 4: *number of replicates passing filters for each metabolome sample*

Supplementary Table 5: *Table of metabolite normalized abundance per sample, Z-scaled across the whole dataset\*

Supplementary Table 6: *List of enriched metabolites per seed compartment, from pairwise Dunn test on BH-adjusted kruskal test p-values*

Supplementary Table 7: *Table of metabolite normalized abundance per sample, Z-scaled per organ*

Supplementary Table 8: *within-compartment PERMANOVA results*

Supplementary Table 9: *OPLS model quality assessed by R²Y and Q²Y averaged across permutations, LOVO cross-validation, and permutation testing*

Supplementary Table 10: *List of candidate metabolites associated with t50germ or FEP*

Supplementary Table 11: *Results of distance-based redundancy analysis (db-RDA) testing the association between seed-associated microbial community composition and variety-level germination and emergence variables*

Supplementary Table 12: *Individual ASV associations with seed germination and emergence traits across four statistical methods*

Supplementary Table 13: *Composition of microbial community inter-kingdom network*

Supplementary Table 14: *Inter-kingdom network assortativity*

## Abbreviations

HR-axis: hypocotyl-radicle axis
FEP: final emergence percentage
FGP: final germination percentage
t50germ: time to reach 50% of final germination.

