## Supplementary Figures for "Identification of Candidate Seed Metabolites and Microbiota Members associated with Germination and Emergence in Common Bean"

**Institutional addresses**
1: Institut Agro, INRAE, IRHS, SFR QUASAV, Université d'Angers, F-49000 Angers, France
2: GEVES, Station Nationale d'Essais de Semences, F-49071 Beaucouzé, France
3: FNAMS, Impasse de verger, 49800 Brain sur l'Authion, France
4: Ecophysiologie et Génomique Fonctionnelle de la Vigne, University of Bordeaux, Bordeaux Science Agro, INRAE, ISVV, Villenave-d'Ornon, France

**Correspondence**

Figure S1: **Supplementary data on experimental design**
Figure S2: **Supplementary data on Germination and Emergence**
Figure S3: **Supplementary data on Morphological characteristics**
Figure S4: **Supplementary data on Metabolomic characteristics**
Figure S5: **Supplementary data on Seed Microbiota characteristics**

#### Experimental design

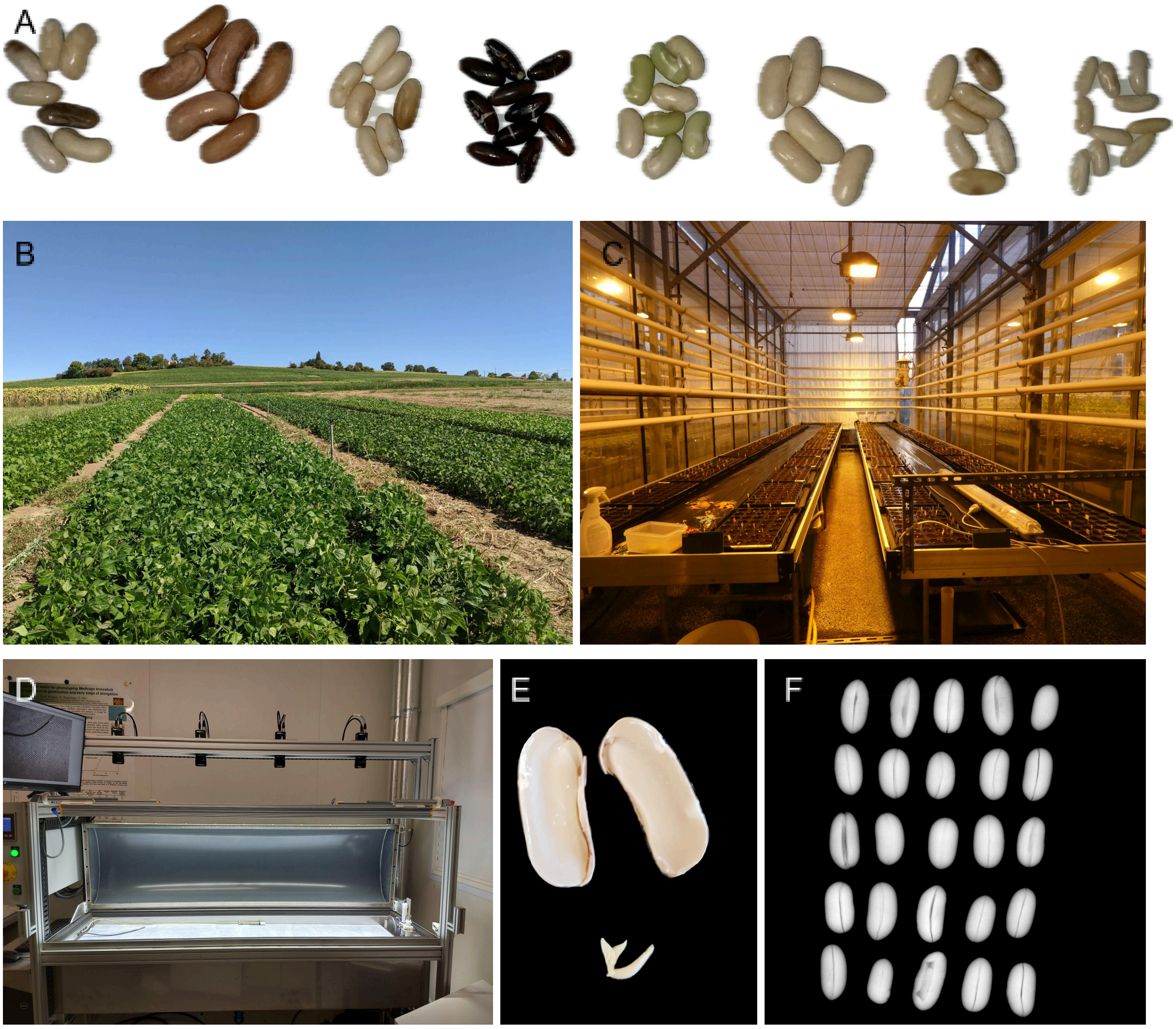


Figure S1: **Supplementary data on experimental design**.

A: Selected common bean varieties. From left to right, top to bottom: Caprice, Contender, Deezer, Facila, Flavert, Linex, Vezer, Vanilla

B: Field assay. Each variety was sown in a 25 x 2.8 m rectangle in the same field at a density of 30 plants per $m^2$, with space between each line and row.

C: View of the greenhouse setup for the emergence assay. Seeds were sown in 40-well trays, and grown in the greenhouse for 7 days.

D: Germination bench used for the germination assays.

E: Dissection of a mature seed into organs for metabolomic analyses.

F: Seed 2D X-ray radiography image as used for the morphological characterisation (here of seeds from variety Caprice).

#### Germination & Emergence

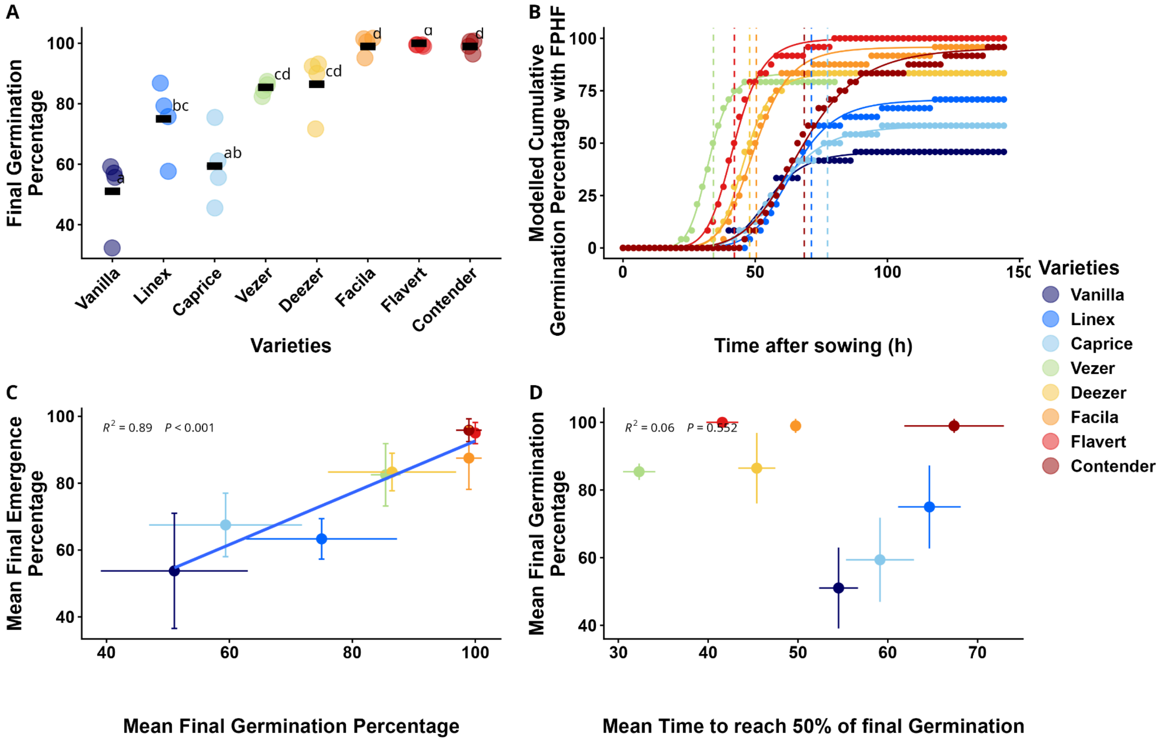


Figure S2: **Supplementary data on Germination and Emergence**

A: **Final germination percentage** for a germination assay on pleated paper, on Jacobsen germinators, at 20°C, 144 h (4*24 seeds). Different letters indicate significant differences (P < 0.05, one-way ANOVA, post-hoc pairwise comparison of adjusted means). Black rectangles: variety mean FEP.

B: **Modeled cumulative germination percentage through time with a Four Parameters Hill Function** (FPHF), from the same germination assay. Curves represent the modeled data, while dots indicate the number of raw data points at given coordinates. Dashed vertical lines indicate the modeled mean time for 50% of final germinated seeds to germinate (mean t50germ).

C: **Correlation of the eight varieties mean final germination percentage** (mean FGP). **with mean time for 50% of final germinated seeds to germinate** (mean t50germ).

D: **Correlation of the eight varieties mean final germination percentage** (mean FGP) **with mean final emergence percentage** (mean FEP)

In C and D, error bars represent SD. Variety colors reflect FEP ranking and are consistent across figures.

#### Morphological characteristics

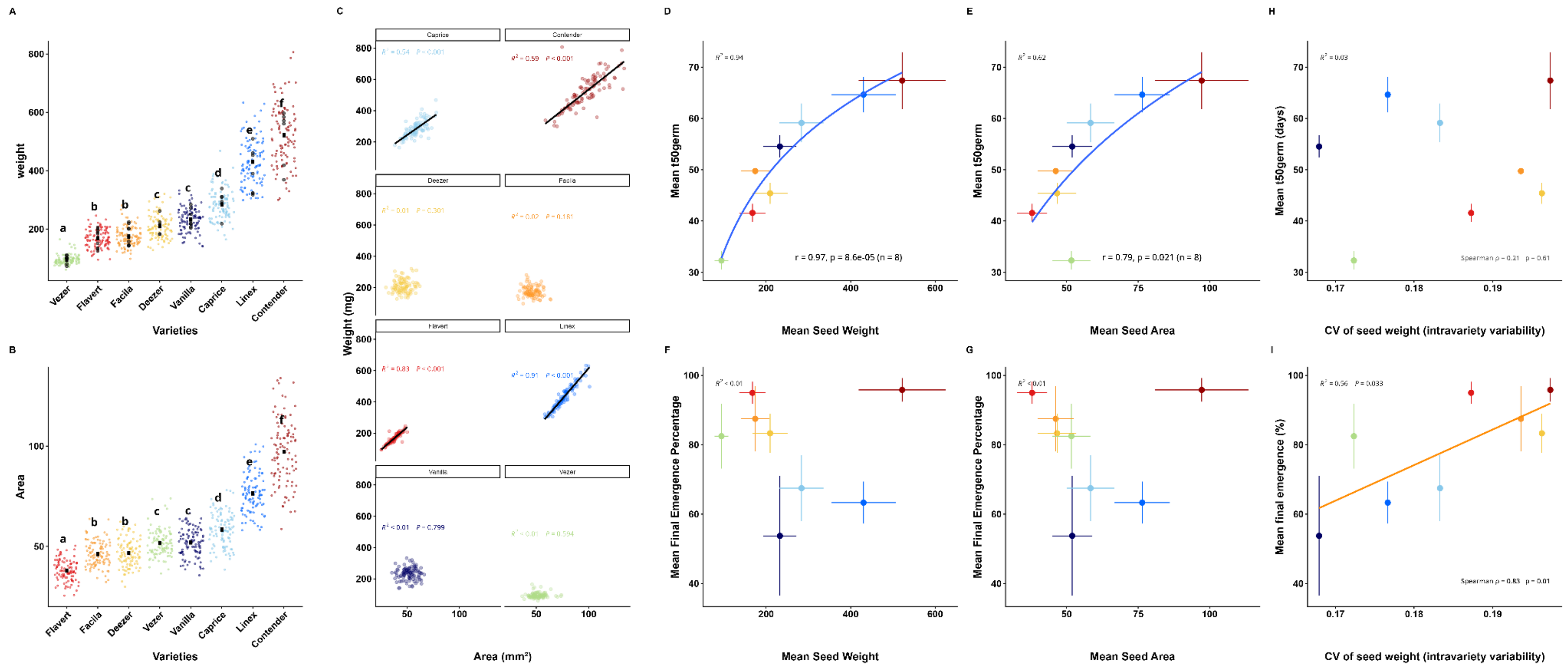


Figure S3: **Supplementary data on Morphological characteristics**

A-B: **Weight (A) or area (B) scatterplot of individual seeds**. Each dot represents one seed n=100). Different letters indicate significantly different mean weight or area (P < 0.05, one-way ANOVA, post-hoc pairwise comparison of adjusted means). (A): dark grey dots represent the additional seeds chosen for metabolomic analyses only.

C: **Weight-Area correlations of individual seeds**. Each dot represents one seed. One color is attributed to each variety and is consistent across panels. Correlation coefficients are written in each by-variety subpanel. Robust regression lines are shown in black.

D-G: **Correlation of average seed weight** (D, F) **or average seed area** (E, G) **of the eight varieties to mean time for 50% of final germinated seeds (mean t50germ)** (D, E) or **mean final emergence percentage** (F, G). Blue curves show logarithmic regression.

H-I: **Correlation of average coefficient of variation of seed weight (cv-weight) of the eight varieties to mean time for 50% of final germinated seeds (mean t50germ)** (H) or **mean final emergence percentage** (I). In D-I, bars represent standard deviation.

#### Metabolomics

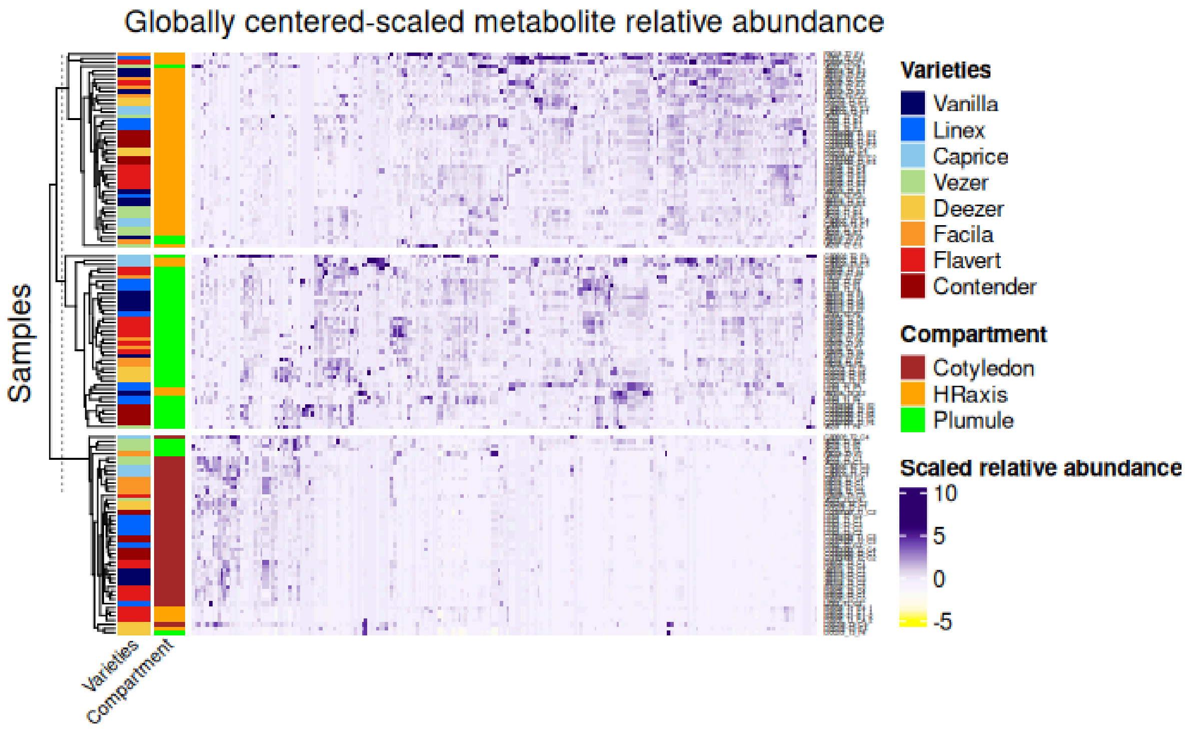


Figure S4. **Heatmap of the relative metabolic abundance of the samples**, clustered in three blocs, with samples in rows (38 cotyledon, 53 HR-axis, and 46 plumule samples) and metabolites in columns (n=221). Colors on the left-hand side represent sample variety (first column) and sample organ (second column). Metabolite abundance was centered-scaled on the whole three-organs dataset.

### Microbiota

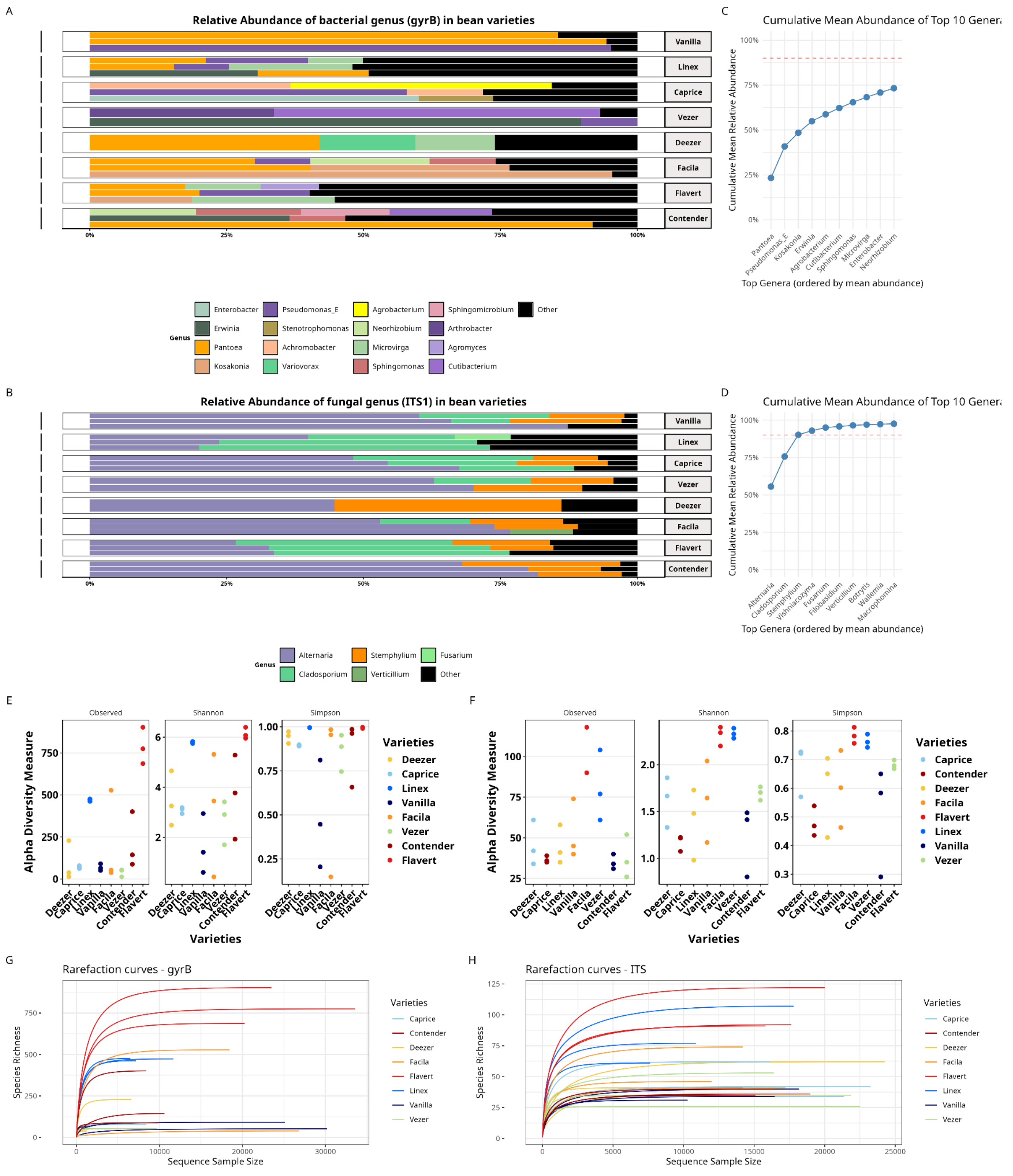


Figure S5: **Supplementary data on seed microbiota characteristics** (n=1-3 samples passing the filters).

A-B: **Relative abundance** of bacterial (C) or fungal (D) taxa, grouped at the genus level. ASVs were filtered on predominance (at least two samples), total count (> 50 across all samples). Genus with relative abundance below 1% were gathered as “Other”. C-D: **Cumulative relative abundance** of top 10 most abundant bacterial (E) or fungal (F) genera. E-F: **Alpha diversity indices** for bacterial (A, *gyrB* marker) or fungal (B, ITS1 marker) communities. G-H: **Rarefaction curves** for bacterial (A, *gyrB* marker) or fungal (B, ITS1 marker) communities.
