## Supplementary Note for "Identification of Candidate Seed Metabolites and Microbiota Members associated with Germination and Emergence in Common Bean"

### Statistical analyses of microbial community composition

**𝛼-diversity**, the microbial diversity measured within a sample, was estimated on the rarefied datasets excluding two bacterial samples with fewer than 1,000 reads (leaving n = 2 for Deezer and Vezer) with richness and Shannon index (that estimates the taxa diversity in a sample taking into account both abundance and evenness). Differences in alpha-diversity among seed varieties were tested using Kruskal-Wallis tests, followed by Dunn's post-hoc pairwise comparisons with Benjamini-Hochberg correction for multiple testing.

**𝛽-diversity**, the microbial diversity compared between samples, was analyzed on unrarefied data filtered by prevalence of at least 5% of samples (454 bacterial and 132 fungal ASVs), then transformed with robust centered-log ratio with *microbiome* (v.1.34.0, (Lahti and Shetty 2017)) which handles zero values without requiring prior imputation. Euclidean distances were computed using *stats::dist*. Community composition was visualized using Principal Coordinates Analysis (PCoA) implemented in the ape R package. The effect of variety on community composition was tested using permutational multivariate analysis of variance (PERMANOVA) with 999 permutations, as implemented in *vegan::adonis2* (v.2.7.3, (Oksanen et al. 2026)). Homogeneity of multivariate dispersions among varieties was assessed prior to PERMANOVA using *vegan::betadisper* followed by a permutation test, to ensure that significant PERMANOVA results reflect differences in community position rather than dispersion. Pairwise post-hoc comparisons between varieties were performed using *pairwiseAdonis::pairwise.adonis2* with Benjamini-Hochberg correction for multiple testing.

Taxonomic composition of seed-associated microbial communities was plotted at genus-level on non-rarefied read counts. ASVs were first filtered by prevalence of at least 5% with at least 50 counts across all samples, then aggregated at genus level. The mean of per-sample relative abundance was computed, and genera of below 1% maximum relative abundance were grouped as “Other” for stacked bar plots.

The taxonomic and occurrence information were used to produce a stacked bar plot of the most predominant genus from taxa with relative abundance superior to 0.1%.

### Seed microbiota composition is variety-dependent for fungi, but not for bacteria

Median Good's coverage after rarefaction was 99.99% for bacterial communities (minimum 99.88%) and 99.97% for fungal communities. Observed ASV richness per sample ranged from 13 to 903 for bacterial communities (median 83.5) and from 26 to 118 for fungal communities (median 41.5), with variety means spanning 40–788 ASVs (gyrB) and 35–99 ASVs (ITS). Flavert and Linex displayed the highest observed richness and Shannon diversity for both bacterial and fungal communities (**Figure S5.E-F**). Both indices differed significantly among varieties for both markers (permutation Kruskal–Wallis, 10⁵ resamples: ITS, χ² = 15.87, p = 0.004 for richness and χ² = 16.73, p = 0.002 for Shannon; gyrB, χ² = 15.54, p = 0.006 and χ² = 15.59, p = 0.006; all Benjamini–Hochberg-adjusted p = 0.006). In Dunn post-hoc tests, only the Flavert–Vanilla contrast for bacterial Shannon diversity remained significant after correction (adjusted p = 0.042). This value equals the smallest adjusted p attainable with three replicates across 28 pairwise comparisons, so only contrasts showing complete rank separation could be detected at this level of replication. All remaining contrasts at adjusted p < 0.10 likewise opposed Flavert or Linex to the least diverse varieties (Contender, Deezer, Vanilla, Vezer).

β-diversity analysis with PCoA of rCLR-transformed community profiles revealed contrasting patterns between bacterial and fungal communities (**Figure 5A-B**). For bacterial communities (*gyrB*), the first two principal coordinates explained 54.3% and 26.9% of total variance, respectively. PERMANOVA indicated no significant effect of variety on bacterial community composition (R² = 0.37, F = 1.09, p = 0.4), and multivariate dispersions were homogeneous among varieties (betadisper, p = 0.397), suggesting that seed variety did not shape bacterial community structure. For fungal communities (ITS), the first two principal coordinates captured 57.7% and 39.7% of total variance, accounting for 97.4% of total variance and indicating a strong two-dimensional structure. PERMANOVA revealed a highly significant effect of variety on fungal community composition (R² = 0.80, F = 7.65, p = 0.001), with homogeneous dispersions among varieties (betadisper, p = 0.25). Given the low number of replicates per variety (n = 2-3), pairwise post-hoc comparisons did not identify any individually significant variety pairs after Benjamini-Hochberg correction.
